# Evolution of salivary glue genes in *Drosophila* species

**DOI:** 10.1101/359190

**Authors:** Jean-Luc Da Lage, Gregg W. C. Thomas, Magalie Bonneau, Virginie Courtier-Orgogozo

## Abstract

**Background:** At the very end of the larval stage Drosophila expectorate a glue secreted by their salivary glands to attach themselves to a substrate while pupariating. The glue is a mixture of apparently unrelated proteins, some of which are highly glycosylated and possess internal repeats. Because species adhere to distinct substrates (i.e. leaves, wood, rotten fruits), glue genes are expected to evolve rapidly.

**Results:** We used available genome sequences and PCR-sequencing of regions of interest to investigate the glue genes in 20 *Drosophila* species. We discovered a new gene in addition to the seven glue genes annotated in *D. melanogaster.* We also identified a phase 1 intron at a conserved position present in five of the eight glue genes of *D. melanogaster*, suggesting a common origin for those glue genes. A slightly significant rate of gene turnover was inferred. Both the number of repeats and the repeat sequence were found to diverge rapidly, even between closely related species. We also detected high repeat number variation at the intrapopulation level in *D. melanogaster*.

**Conclusion:** Most conspicuous signs of accelerated evolution are found in the repeat regions of several glue genes.

## Background

Animals interact with their environment (viruses, bacteria, food, chemicals, conspecifics, etc.) in many different ways, particularly through their immune and sensory systems. As animals adapt to new places, the way they interact with their environment is expected to change. Accordingly, the gene families that have been shown to exhibit accelerated rates of gene gain and loss in several animal groups are mostly genes that mediate the interactions with the environment: immune defense, stress response, metabolism, cell signaling, reproduction and chemoreception [1]. Rapid changes in gene copy number can lead to fast phenotypic changes via gene deletion and can provide raw material for genes with new functions via gene duplication [2]; [3]. Rapid turnover of genes within a gene family has also been shown to correlate with fast evolution at the sequence level [4, 5].

Here we focus on the Salivary gland secretion (*Sgs*) genes, a functional group that mediates the physical interaction of flies in the genus *Drosophila* with an external substrate during metamorphosis. The *Sgs* genes encode proteins that make up the glue produced by Drosophila larvae that serves to attach the animal to a surface where it can undergo metamorphosis. In *D. melanogaster*, the glue is composed of several salivary gland secretion proteins which accumulate in the salivary glands of late third instar larvae [6]. As the puparium forms, the bloated salivary glands release their contents through the mouth. This secretion then hardens within seconds of contact with the air and becomes a glue which firmly attaches the pupa to the substrate. Metamorphosis is a critical stage of Drosophila development [7] during which the animal is vulnerable and motionless. In Drosophilids pupae are generally attached to a substrate until the imago leaves the puparium. It is critical for the pupa to be firmly attached in order not to be moved away by some external event (i.e. rain or wind). Furthermore, for the emerging adult to be able to hold on the external substrate and thus get out of the pupal case, it is necessary for the pupa to adhere to a substrate, whether dry or wet. When the pupal case freely moves and is not attached, adults are unable to hatch and eventually die (J. R. David, personal communication).

Pupation sites of *Drosophila* species in nature have not been extensively characterized, but a large diversity of pupation sites have been found. In the wild, *D. melanogaster* pupae have been found adhered to wood, fixed to grape stalks, attached to the dry parts of various rotten fruits, or adhered to one another on the land beneath grape stalks (J. R. David, personal communication, [8-10]). *D. mauritiana* pupae may be found on the surface of decaying *Pandanus* fruit, which is hard and lignous (D. Legrand, personal communication). Many Hawaiian *Drosophila* species pupariate several inches deep in the soil [11]. Some other *Drosophila* species, such as *D. sechellia, D. simulans, and the invasive D. suzukii*, appear to pupariate directly within the wet rotten part of fruits (J. David, personal communication, [12]).

Given the diversity of pupation sites, we hypothesized that they would require distinct types of glue and therefore that *Sgs* genes might evolve rapidly among the *Drosophila* genus. The glue genes have long been an important model for the regulation of gene expression since it was discovered in the 1970-80s that genes for proteins contained in salivary secretions were found to correlate with the chromosomal location of major puffs (13-16). This led to the discovery that, on an acid-urea electrophoresis gel, the salivary glue was resolved into five major bands, numbered from 1 to 5 in order of increasing electrophoretic mobility [16, 17], Band 2, which was variable and detected in many other tissues, was considered to be a tissue contamination rather than a true glue protein [16] and from this seven glue genes were eventually identified, and their nucleotide sequences are now well characterized: *Sgs1* (band 1, *CG3047*, 2L), *Sgs3* (band 3, *CG11720*, 3L), *Sgs4* (band 4, *CG12181*, X), *Sgs5* (band 5, *CG7596*, 3R), *Sgs7* (*CG18087*, 3L), and *Sgs8* (*CG6132*, 3L) and *Eig71Ee* (also named *geneVII I71-7* or *gp150, CG7604*, 3L) [14, 18-27]. *Eig71Ee*, located at position 71E, is not only expressed in salivary glands but also in hemocytes and in the gut, where it appears to be involved in immunity and clotting [28-30].

A sixth electrophoretic band migrating slightly slower than the Sgs3 protein was also detected in a few *D. melanogaster* lines [17, 31, 32]. The nucleotide sequence of the corresponding gene, *Sgs6*, remains unknown but cytogenetic and genetic mapping indicates that *Sgs6* is located in region 71C3-4 and differs from *Eig71Ee* [14, 28, 32].

The three genes *Sgs3*, *Sgs7* and *Sgs8* form a tightly linked cluster on the 3L chromosomal arm at position 68C [33, 34]. All glue genes were found to start with a signal peptide. The largest glue genes, *Sgs1*, *Sgs3* and *Sgs4* and *Eig71Ee* were shown to harbor numerous internal repeats of amino acid motifs, rich in proline, threonine and serine [19, 25, 29, 35]. Molecular studies showed that the number of internal repeats was variable between strains in Sgs3 [36], and Sgs4 [35]. Additionally, consistent with missing protein bands, a few laboratory strains were inferred to carry loss-of-function mutations in *Sgs4* [6, 16, 35, 37], *Sgs5* [27] and *Sgs6* [17, 31, 32].

In the present study, we characterize the diversity and evolution of the *Sgs* genes within the *Drosophila* genus. We inferred loss and gain of glue genes and we investigated repeat number variation and sequence repeat diversity across 19 species and across paralogs.

## Results

We used the six *Sgs* genes and *Eig71Ee* annotated in *D. melanogaster* as BLAST queries to identify their putative homologs in 19 other *Drosophila* species (Table 1). The homologs are summarized in Figure 1 and Table 2. In *D. melanogaster*, the glue genes are “extremely highly” or “very highly” expressed in late larval salivary glands according to the RNAseq data in Flybase. But transcript data that would be useful for annotating the genes were not available for all species, probably because the expression window of the glue genes (late third larval instar and only in salivary glands) is narrow [6]. The organization of the *Sgs* genes was found to be generally conserved across the *Drosophila* species we investigated (Fig. 1). Proper identification of each ortholog was based on sequence similarity and, when possible, synteny. We describe below our findings for each category of *Sgs* genes.

**Figure 1.**
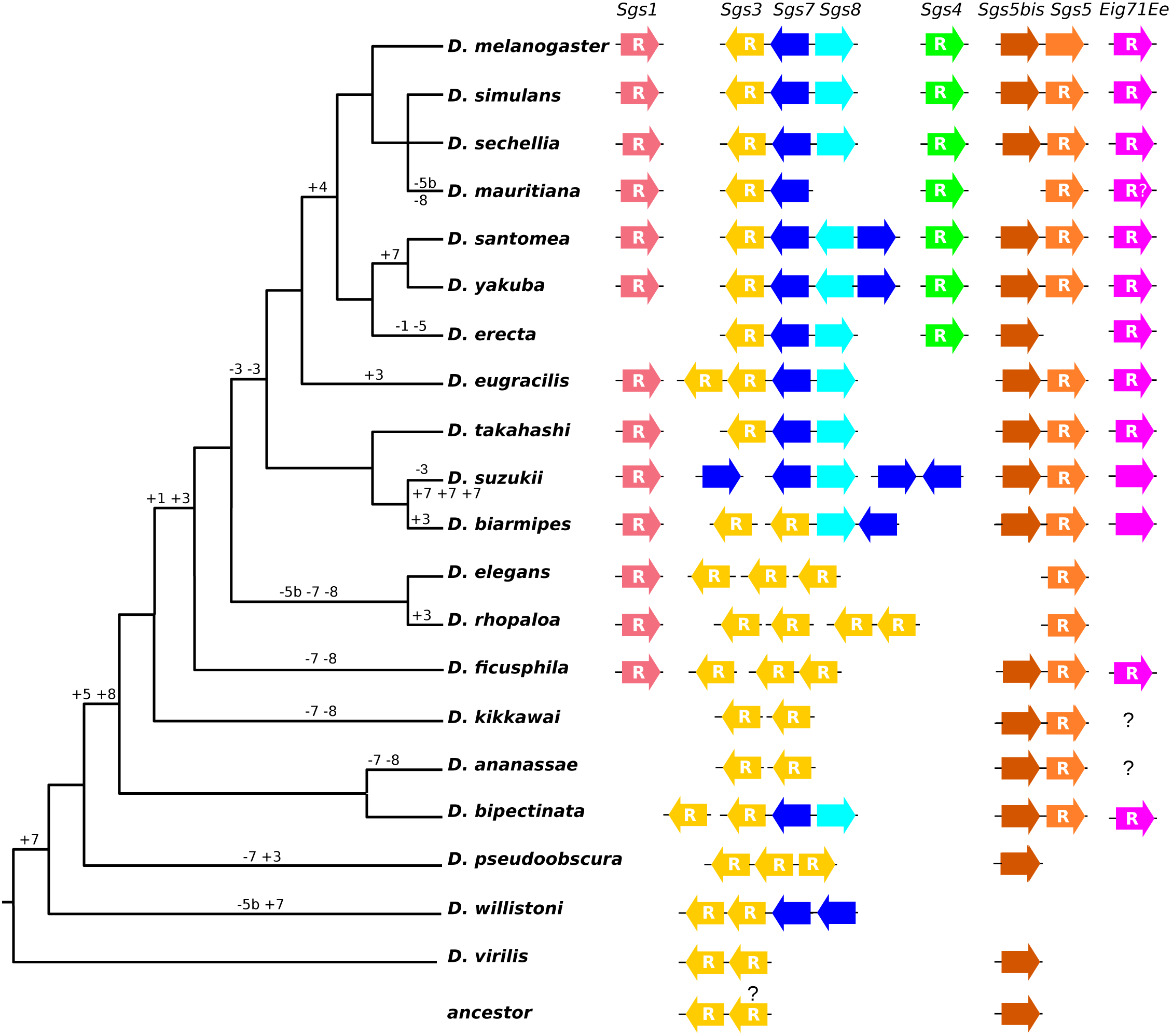
Schematic species tree showing glue gene distribution and the most parsimonious scenario for gene gains and losses. Gains are indicated by “+” and losses by “-”. Numbers correspond to the glue gene name (eg. “3” for Sgs3). An inferred distribution of glue genes in the last common ancestor is shown at the bottom. The tree is from Thomas, G.W.C. and Hahn M.W. (2017) http://dx.doi.org/10.6084/m9.figshare.5450602. Pink is for *Sgs1*, yellow is for *Sgs3*, dark blue is for *Sgs7*, light blue is for *Sgs8*, green is for *Sgs4*, orange is for *Sgs5-5bis*, purple is for *Eig71Ee*. Along with each species is a schematic representation of the organization of the glue gene cluster, with relative position and orientation for the species with confirmed synteny information. Gene sizes and distances are not to scale. “R” means that internal repeats are present. “R?” means that no clear repeats were identified. In *D. pseudoobscura*, the relative orientation of the three clustered *Sgs3*-like sequences GA25425, GA23426, GA23878 suggested that GA23426 could be orthologous to *Sgs3* (it is inside an intron of GA11155, homologue of Mob2, which is close to *Sgs3* in *D. melanogaster*), GA23425 to *Sgs7* and GA23878 to *Sgs8*. The last two had more similar sequences compared to GA23426, including the repeat region. Furthermore, the latter was neighbor to GA20420, a homologue of *chrb-PC*, a gene adjacent to *Sgs8* in *D. melanogaster*.

**Table 1:** List of species and databases used in this study.

| Species | Database | Version | URL | Date of access | reference |
| --- | --- | --- | --- | --- | --- |
| *melanogaster* | FlyBase | FB2015_02 | flybase.org | 06/2016 | [58] |
| *simulans* | FlyBase | FB2015_02 | flybase.org | 02/2017 | [58] |
| *sechellia* | FlyBase | FB2015_02 | flybase.org | 02/2017 | [58] |
| *mauritiana* |  | v1.0 | www.popoolation.at/mauritiana_genome/ | 12/2016 | [61] |
| *yakuba* | FlyBase | FB2015_02 | flybase.org | 02/2017 | [58] |
| *santomea* |  | v1.0 | genomics.princeton.edu/AndolfattoLab/Dsantomea_genome.html | 11/2016 | [60] |
| *erecta* | FlyBase | FB2015_02 | flybase.org | 02/2017 | [58] |
| *takahashii* | FlyBase | FB2015_02 | flybase.org | 02/2017 | [58] |
| *ficuspila* | FlyBase | FB2015_02 | flybase.org | 02/2017 | [58] |
| *biarmipes* | FlyBase | FB2015_02 | flybase.org | 02/2017 | [58] |
| *suzukii* | SpottingWingFlybase | v1 | http://spottedwingflybase.org/ | 02/2017 | [59] |
| *eugracilis* | FlyBase | FB2015_02 | flybase.org | 02/2017 | [58] |
| *elegans* | FlyBase | FB2015_02 | flybase.org | 02/2017 | [58] |
| *rhopaloa* | FlyBase | FB2015_02 | flybase.org | 02/2017 | [58] |
| *kikkawai* | FlyBase | FB2015_02 | flybase.org | 02/2017 | [58] |
| *ananassae* | FlyBase | FB2015_02 | flybase.org | 02/2017 | [58] |
| *bipectinata* | FlyBase | FB2015_02 | flybase.org | 02/2017 | [58] |
| *willistoni* | FlyBase | FB2015_02 | flybase.org | 02/2017 | [58] |

**Table 2:** Genomic coordinates of the glue genes in 20 *Drosophila* species. * indicates annotations and coordinates of the *Sgs5bis* gene; “M” indicates that part of the coding sequence was inferred manually by sequencing of PCR amplicons of relevant regions; “no” means that the gene sequence was not found by BLAST searches; Nterm and Cterm mean N-terminal and C-terminal region, respectively. **: this contig probably contains two paralogs of *Sgs3* with uncertain sequences.

| Species | *Sgs1* | *Sgs3* | *Sgs4* | *Sgs5 Sgs5bis\** | *Sgs7* | *Sgs8* | *Eig71Ee* |
| --- | --- | --- | --- | --- | --- | --- | --- |
| *D. melanogaster* | CG3047 | CG11720 | CG12181 | CG7596<br>CG7587* | CG18087 | CG6132 | CG7604 |
| *D. simulans* | GB:CM002910<br>4752550- 4754973 | Dsim\GD14311 | Dsim\GD1663<br>7 | Dsim\GD19170<br>Dsim\GD19169* | Dsim\GD17634 | Dsim\GD28639 | Dsim\GD12546 |
| *D. sechellia* | Dsec\GM18501<br>(M) | Dsec\GM25279 (M) | GB:CH480825<br>2852711-2853386 (M) | Dsec\GM15245<br>Dsec\GM15244* | Dsec\GM25278 | Dsec\GM24748 | NW_001999689<br>7761215-7759941 |
| *D. mauritiana* | 2L : 4721427-4722731 | 3L : 11002313-11003109 | X : 2864998-2865616 (M) | 3R : 7695225-7694660<br>relictual *Sgs5bis*<br>3R : 7696600-7695629 | 3L : 10999955-11000249 | no | 3L : 15018149-15017249 |
| *D. yakuba* | NT_167062<br>10588365-10585585 | Dyak\Sgs3 | Dyak\GE28681 | Dyak\GE25481<br>Dyak\GE25480* | Dyak\GE20214<br>Dyak\GE21218 | Dyak\Sgs8 | Dyak\GE19823 |
| *D. santomea* | 2L : 10595909-10588129 | 3L : 11541799-11542678 (M) | X : 5242740-5241688 (M) | 3R : 1975190-1975883<br>3R : 1974195-1974756* | 3L : 11539572-11539861<br>3L : 11536774-11536485 | 3L : 11537383-11537681 | 3L : 18202978-18201736 |
| *D. erecta* | no | Dere\Sgs3 | Dere\GG27095 | no *Sgs5*<br>Dere\GG22329* | Dere\GG13918 | Dere\Sgs8 | Dere\GG13528 |
| *D. eugracilis* | AFPQ02004874<br>817906- 819883 | KB465257<br>3401691- 3402412<br>3385186-3386300 | no | KB464468<br>62658- 63338<br>61657-62202* | KB465257<br>3378701- 3378995 | KB465257<br>3378110- 3377822 | KB464880<br>383836-382228<br>(XM_017230731) |
| *D. takahashii* | KB461520<br>248469-250276 | KB460792<br>317161-317949 | no | KB461611<br>188299-187637<br>189545-188599* | KB461234<br>120246-120467 | KB461234<br>119117-118896 | XM_017142344 |
| *D. ficusphila* | KB457325<br>1315471-1313145 | KB457563<br>3180441-3179541 | no | KB457381<br>2059719- 2058971 | no | no | KB457515<br>1660700-1661809 |
|  |  | KB457373<br>332100-331262<br>3199436-3198351 |  | 2061615-2060148* |  |  | (XM_017197540) |
| *D. biarmipes* | KB462641<br>1521394-1523538 | KB462590<br>1536842-1537624 (M)<br>KB462646<br>54238-53374 (M) | no | KB462814<br>8082338-8083047<br>8081336-8081891* | KB462646<br>76095-75801 | KB462646<br>77216-77501 | KB462754<br>733209-734564 |
| *D. suzukii* | KI419149<br>6645021-6638237 | no | no | KI420542<br>10372-9639<br>11441-10912* | KI419359<br>22757-22464<br>KI420769<br>54293-54584<br>KI420610<br>25121-25412<br>55385-55094 | KI420769<br>53260-52976 | XM_017082231 |
| *D. elegans* | KB458429<br>2603084- 2605600 | KB458268<br>2467758- 2468497<br>KB458387<br>820622-819957<br>KB458387<br>18429-17499 | no | KB458458<br>2864199- 2863401<br>no Sgs5bis | no | no | no |
| *D. rhopalosa* | KB450401 (Nterm)<br>KB452165 (Cterm) | KB450817<br>117692-118515<br>KB452471<br>215593-216424<br>KB451944** | no | KB451039<br>15186-16018<br>no Sgs5bis | no | no | no |
| *D. kikkawai* | no | KB459615<br>1331679-1331220<br>KB459522<br>291906-292542 | no | KB459676<br>1112222-1111011<br>1113233-1112671* | no | no | KB459876<br>1106397-1107027<br>(Nterm) |
| *D. ananassae* | no | NW_001939300<br>3959435-3957637<br>NW_001939293<br>5806878-5808646 | no | NW_001939291<br>17741832-17741201<br>17742892-17742284* | no | no | GF10382(Nterm):<br>NW_001939293<br>11506744-11507112 |
| *D. bipectinata* | no | KB464001<br>557673-558039<br>KB464098<br>1120437-1121198 | no | KB464382<br>185749-186362<br>184743-185354* | KB464098<br>1109828-1110127 | KB464098<br>1109077-1108802 | KB464259<br>2466431-2466234<br>(ortholog of<br>GF10382) |
| *D.* | no | GA23425, GA23426, | no | no Sgs5 | no | no | no |
| *pseudoobscura* |  | GA23878 |  | Dpse\GA20459 * |  |  |  |
| *D. willistoni* | no | NW_002032853<br>3296683-3295766<br>NW_002032860<br>11643758-11641972 | no | no | NW_002032853<br>2792051-2792347<br>2793811-2794107 | no | no |
| *D. virilis* | no | NW_002014431<br>6839085-6838999<br>(GJ27025)<br>6841799-6840888(GJ26085) | no | no Sgs5<br>NW_002014424<br>14511530-14512000*(GJ24445) | no | no | no |

### Gains and losses of Sgs5 *genes*

We found that *Sgs5* had a tandem paralog in *D. melanogaster*, located ca. 300 bp upstream of *Sgs5* (*CG7587*, hereafter named *Sgs5bis*), sharing 46,3 % identity and 66,9 % similarity at the protein level. It is co-expressed with *Sgs5* during late third larval instar in dissected salivary glands, as shown by similar expression profiles on Gbrowse at flybase.org and FlyAtlas at flymine.org. To our knowledge, this paralog has not been mentioned earlier. Both paralogs harbor two introns in all species. The *Sgs5/5bis* pair is widely distributed and therefore probably ancestral to most of the species studied. The occasional loss of either *Sgs5* or *Sgs5bis* occurred at least four times (Fig. 1): 1) loss of *Sgs5bis* in *D. mauritiana*, where a relictual sequence may still be recognized, 2) loss of *Sgs5bis* in *D. elegans,* 3) loss of *Sgs5bis* in *D. rhopaloa*, 4) loss of *Sgs5* in *D. erecta*. These patterns of loss suggest that *Sgs5* and *Sgs5bis* can replace each other functionally. In *D. ananassae,* the orthologous sequence of *Sgs5bis* (formerly Dana\GF19880 in FlyBase release R1.3, with a different intron/exon structure) has been withdrawn from the genome annotation for reasons unknown to us and it is in conserved synteny relative to *D. melanogaster.* In *D. virilis* and *D. pseudoobscura*, a single *Sgs5*/*5bis* gene was identified. A phylogeny of all Sgs5 and Sgs5bis amino acid sequences (Fig. 2) revealed a clear separation in the gene sequences of the two groups, *Sgs5* and *Sgs5bis*. The *D. virilis* gene annotated as *Sgs5* (Dvir\GJ24445) and the *D. pseudoobscura Sgs5/5bis* gene (annotated as uncharacterized protein Dpse\GA20459) were clustered with the *Sgs5bis* genes and shared with most other *Sgs5bis* sequences a motif Gln-Ala-Thr in the signal peptide. This suggests that *D. virilis* and *D. pseudoobscura* possess an ortholog of *Sgs5bis*. If the *Sgs5-Sgs5bis* gene duplication arose after the separation of the *D. virilis* and *D. pseudoobscura* lineages, then the ancestral gene before the duplication was probably *Sgs5bis.*

**Figure 2.**
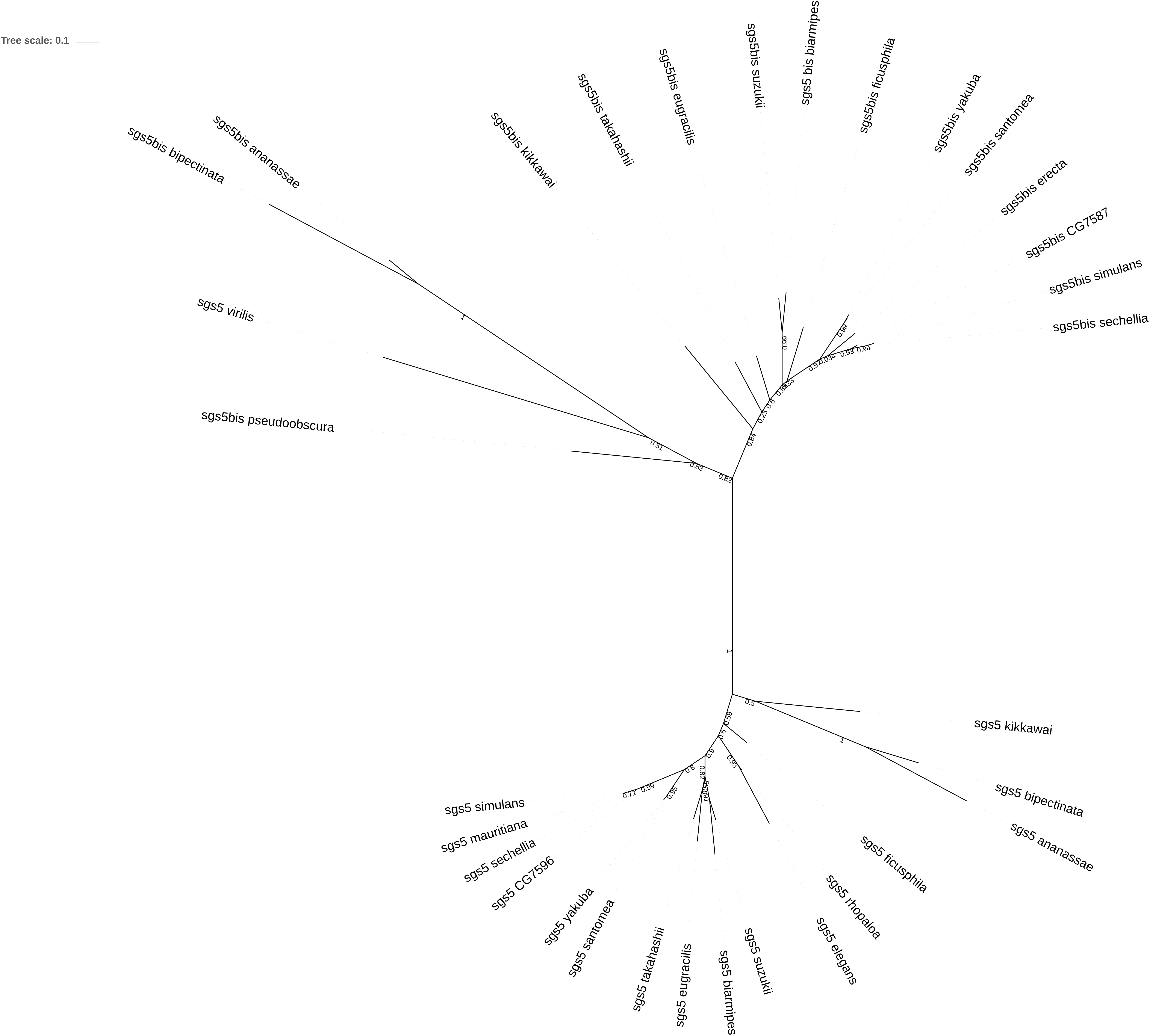
Maximum likelihood (ML) unrooted tree of aligned Sgs5 and Sgs5bis amino acid sequences (repeated parts removed when present). Numbers along branches are the posterior probabilities.

**Figure 3.**
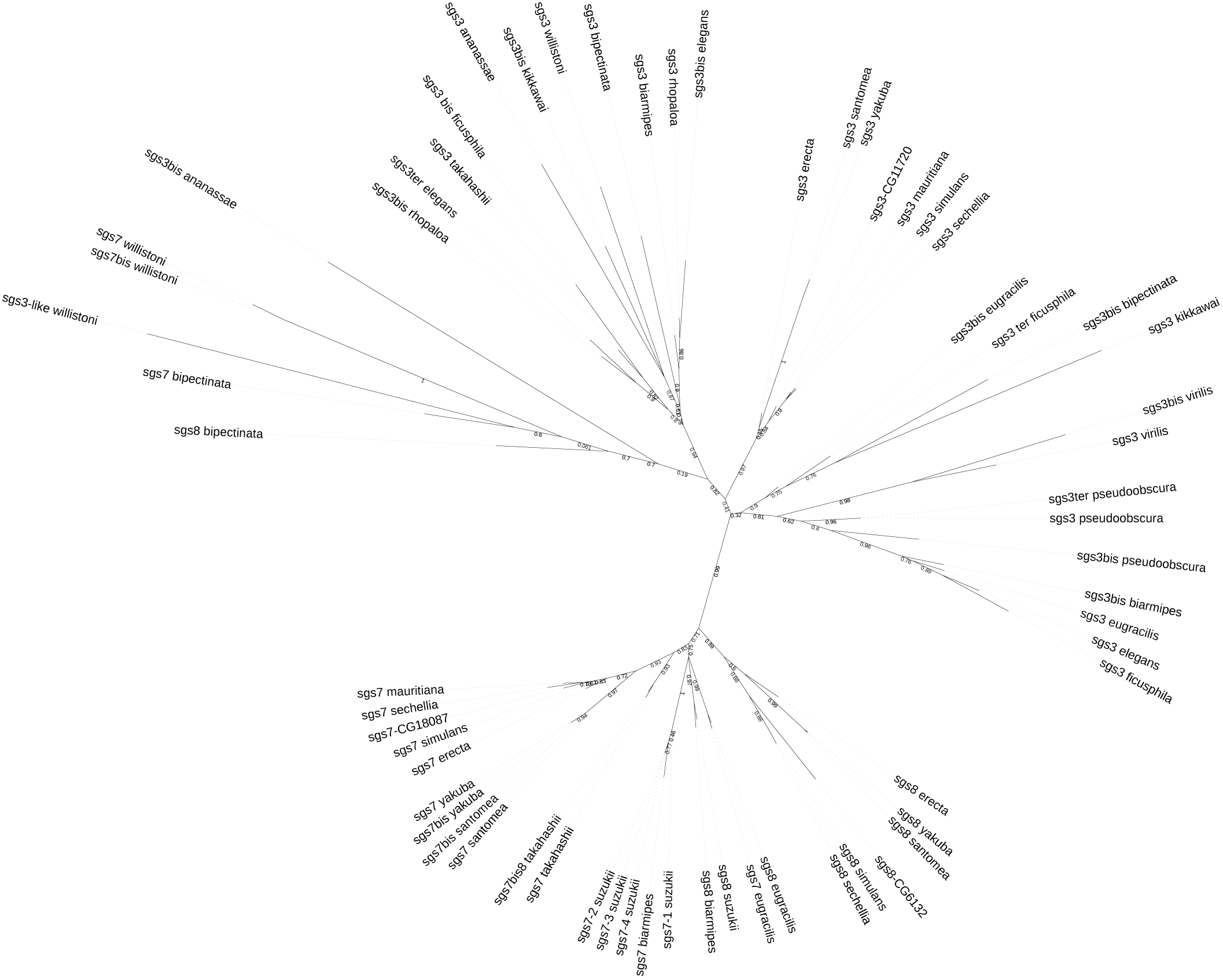
Unrooted ML tree of aligned Sgs3 (repeats removed), Sgs7 and Sgs8 amino acid sequences. Numbers along branches are the posterior probabilities.

### *Gains and losses of* Sgs3, Sgs7, and Sgs8 *genes*

The genes *Sgs3*, *Sgs7* and *Sgs8* form a tight cluster, 4.5 kb long, on the 3L arm in *D. melanogaster* [33] and they share sequence similarities [19] in their N-terminal and C-terminal parts. *Sgs3* contains internal repeats whereas *Sgs7* and *Sgs8* do not. When the internal repeats of *Sgs3* are excluded, the amino acid identity in *D. melanogaster* is 51.3 % between Sgs3 and Sgs7, 48.7 % between Sgs3 and Sgs8, and 46.7 % between Sgs7 and Sgs8. Additionally *Sgs3*, *Sgs7* and *Sgs8* share a phase 1 intron position, interrupting the signal peptide sequence [19]. In the clade *D. yakuba* / *santomea* / *erecta*, *Sgs7* and *Sgs8* are inverted with respect to the *D. melanogaster* arrangement (Fig. 1). In addition, *Sgs7* is duplicated in *D. yakuba* (Dyak\GE20214 and Dyak\GE21218) and *D. santomea* (Fig. 1). The two copies, inverted relative to each other, have only one, nonsynonymous, nucleotide difference. *Sgs8* lies between the two *Sgs7* copies, and has the same orientation as *Sgs3*. In species outside the *D. melanogaster* subgroup, all the *Sgs3*, *Sgs7* and *Sgs8* sequences also have the same intron, with slightly different positions depending on codon indels before the intron. Notably, *D. suzukii* is the only species in our study that has lost *Sgs3. D. suzukii* retained *Sgs8* and underwent an amplification of *Sgs7*, three copies of which are identical. In a number of species, *Sgs7* and *Sgs8* could not be identified (Sgs7 and Sgs8 are small proteins, about 75 amino acids in length). However, when a BLAST search was performed using the *Sgs7* or *Sgs8* sequences of *D. melanogaster*, we retrieved the same target hits as with *Sgs3* (Table 2). In those species, several *Sgs3*-like genes were found instead, i.e. long proteins with internal repeats showing N-terminal and C-terminal parts similar to *Sgs3*. In species with no *Sgs7*, no *Sgs8* and several *Sgs3*-like genes occupying the physical location of *Sgs7* and *Sgs8* (*D. pseudoobscura*, *D. ficusphila*, *D. rhopaloa*, see Fig. 1), it is tempting to infer that the ancestral *Sgs7* and *Sgs8* have gained internal repeats. According to such a hypothesis, at least in some cases, the non-repeated parts of those Sgs3-like protein sequences are expected to cluster with Sgs7/8. To disentangle the relationships among *Sgs3*-7-8 paralogs, we constructed a phylogeny using an alignment of the non-repeated parts of the protein sequences. The tree (Fig. 3), which does not fit well to the species phylogeny, shows a clear separation between *Sgs3/Sgs3*-like and *Sgs7/Sgs8*, except for *D. bipectinata* and *D. willistoni*, whose *Sgs7/Sgs8* sequences are linked to the *Sgs3* branch, with low support. This would rather suggest that those *Sgs7/Sgs8* sequences are old *Sgs3*-like sequences which have lost their internal repeats.

However, the sequence length is far too short to get a reliable tree and we cannot confirm this hypothesis. While it is more parsimonious to infer that there were two ancestral *Sgs3* and that subsequent losses occurred, the tree topology is not accurate enough to confirm this hypothesis.

### *Gains and losses of* Sgs1 *genes*

*Sgs1* was found only in the *melanogaster* subgroup and in the Oriental subgroups, which suggests that it originated in the ancestor of this clade. No *Sgs1* gene was detected in *D. erecta*, providing evidence for a loss of *Sgs1*. The *Sgs1* sequence identified by our BLAST search in the *D. suzukii* genome database (see Materials and Methods) showed many stop codons in the second half of the repeat region and was not annotated as a coding sequence. Based upon the surrounding repeat sequences, we found that inserting a C at position 1829 (from start) would restore the reading frame, translating into a putative 2245 amino acid protein. Our analysis of another genome sequence of *D. suzukii* [38] (contig CAKG01017146) showed that in this second strain there is a C at position 1829 and that Sgs1 is 2245 amino acid long. Since position 1829 lies in the middle of a long repeat-containing region which prevents PCR amplification, we did not try to check experimentally for the missing C in the first *D. suzukii* genome sequence. In all the *Sgs1* genes identified, except in *D. elegans*, an intron was found at the same position and phase as in *Sgs3*, *Sgs7* and *Sgs8*. There is also a loose similarity in the N-terminal and C-terminal parts of Sgs1 and Sgs3 (in *D. melanogaster* about 14% identity between Sgs3 and Sgs1 excluding the repeats). This suggests that *Sgs1* belongs to the same family as *Sgs3/Sgs7/Sgs8* genes.

### *Origin of* Sg4 *and* Eig71Ee *genes*

*Sgs4*, which is intronless, was absent outside the *D*. *melanogaster* subgroup (Fig. 1, Table 2). The origin of *Sgs4* is unknown. We found no similarity with any other sequence in any genome. Some sequence similarity between *Eig71Ee* and *Sgs4* had been reported [29], but is not convincing since it was in the repeat parts, which are of low complexity. *Eig71Ee* was found in all the *D. melanogaster* subgroup species and in some of the so-called Oriental species, where it has been annotated as *mucin2*, or *extensin* in *D. takahashii*, or even, erroneously, *Sgs3* in *D. suzukii*. We also detected the N-terminal parts of it in the *D. ananassae* group; thus making unclear the phylogenetic distribution of the gene (Table 2). More interestingly, we noticed that *Eig71Ee* harbors an intron at the same position as the one of *Sgs3, Sgs7, Sgs8* and *Sgs1*. This result argues for a certain relatedness among those genes. However, using Eig71Ee as a TBLASTN query did not retrieve any hits from any *Sgs* genes and the Eig71Ee amino acid sequence does not align with the Sgs sequences.

### Rate of gene gains and losses in the glue gene families

Our analysis reveals that the seven annotated genes that code for glue proteins can be grouped into three gene families. *Sgs1, Sgs3, Sgs7, Sgs8,* and *Eig71Ee* comprise one of the three families since all of them share a phase 1 intron at the same position, interrupting the signal peptide sequence. *Sgs4* then forms its own family and the *Sgs5* and *5bis* comprise the third family. We used CAFE [39] to reconstruct ancestral copy numbers throughout the Drosophila phylogeny and to test whether these three gene families evolve at an accelerated rate along any Drosophila lineage. For the CAFE analysis *Eig71Ee* was not included due to uncertainties about its presence in some species. We find that the *Sgs4* and *Sgs5-5bis* families do not evolve faster compared to other gene families present in the Drosophila genomes (p=0.58 and p=0.107, respectively; Table S1), however the *Sgs1-3-7-8* family was found to evolve rapidly (p=0.005; Table S1). Overall, this family seems to be prone to duplication and loss (Fig. S1) and we find that this signal for rapid evolution is driven mostly by small changes on many lineages (i.e. a gain or loss of 1 gene) rather than large changes on one or a few particular lineage.

### Characterization of the repeats in glue proteins

Table 3 summarizes the characteristics of the repeated sequences present within *Sgs* genes. Sgs1, Sgs3 and to a lesser extent, Sgs4 and Eig71Ee, are characterized, besides a signal peptide and a conserved C-terminal part, by long repeats often rich in threonine and prone to O-glycosylations. Although *D. melanogaster* Sgs5 protein is devoid of internal repeats (we checked that it is the case in all populations of the PopFly database), in most other species, even in close relatives, repeats are present, mostly pairs Pro-(Glu/Asp). Sgs5 protein length is highly variable across species. In *D. kikkawai*, there is a long additional stretch (127 amino acids) containing 60 % of acidic residues. The paralog Sgs5bis never has repeats. Sgs7 and Sgs8 are much smaller proteins, without any repeats, and are rich in cysteine (12-14 %). The conserved C-terminal parts are about 120 amino acids long in Sgs1, 50 amino acids in Sgs3, 120 amino acids in Sgs4, 115 amino acids in Sgs5/5bis and 135 amino acids in Eig71Ee. The repeats are quite variable in motif, length and number, even between closely related species, so that, most often, glue proteins may be retrieved only based on their conserved C-terminal part. The longest Sgs protein is Sgs1 of *D. suzukii* (2245 aa), which harbors ca. 63 repeats of a 29 amino acid, threonine-rich motif so that the total content of threonine is 40%; in *D. melanogaster*, Sgs1 is also very long (1286 aa) due to 86 repeats of a motif of 10 amino acids, also threonine-rich (46%). The shortest Sgs1 protein is the one of *D. sechellia* (492 aa). In all the species where it exists, Sgs1 is also rich in proline (12-18%). Sgs3 has the same kind of amino acid composition. Repeats can also be quite different between paralogs. For example, in *D. eugracilis*, while the two genes are physically neighbors, Sgs3 has about seven repeats of CAP(T)_9_, whereas Sgs3bis has ca. 65 KPT repeats. In *D. elegans*, the three Sgs3-like proteins also have quite different repeats (Table 3). Sgs4 is richer in proline than in threonine (18% vs. 16% in *D. melanogaster*) and contains 10% cysteine residues.

**Table 3:**
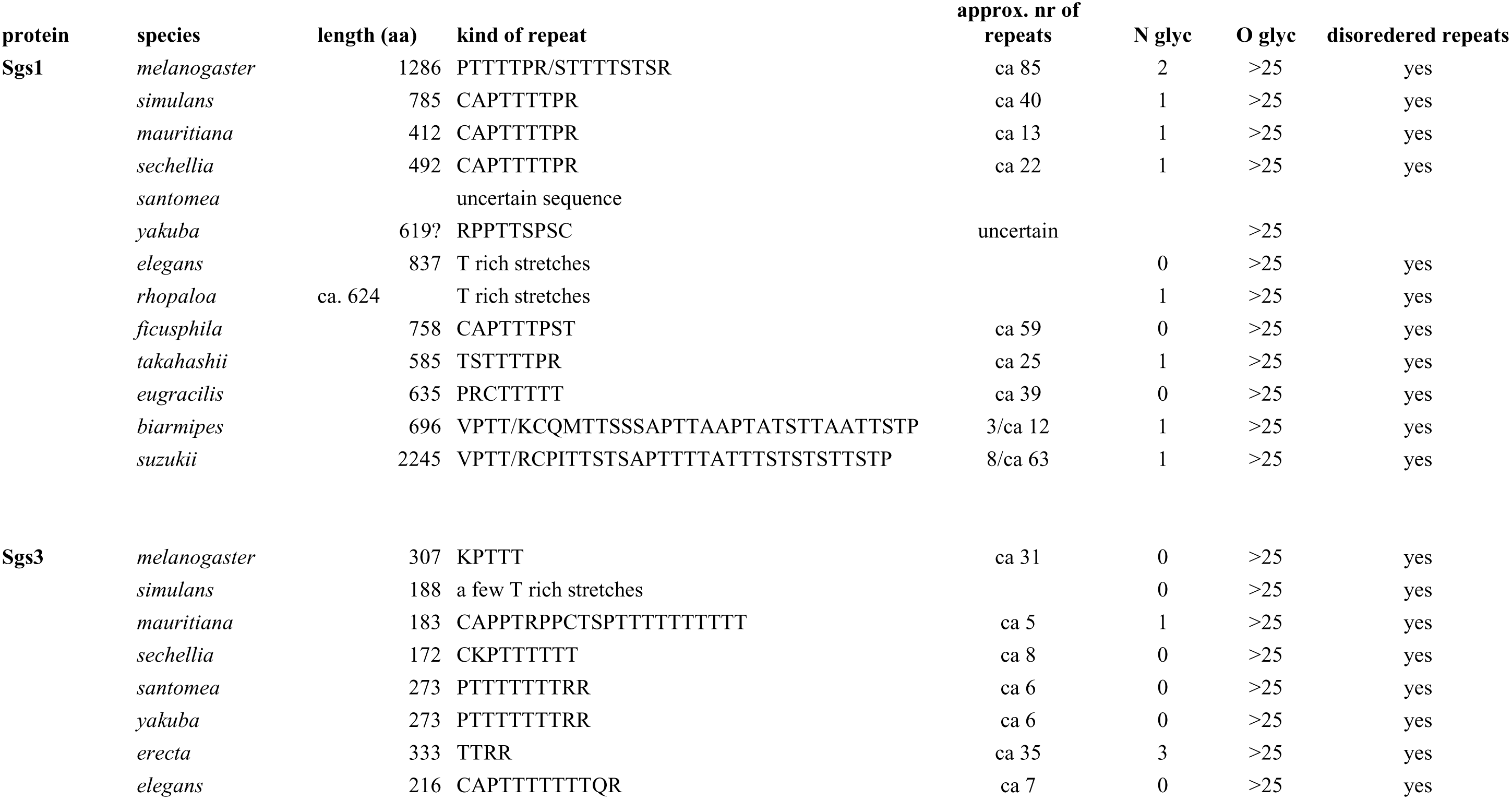
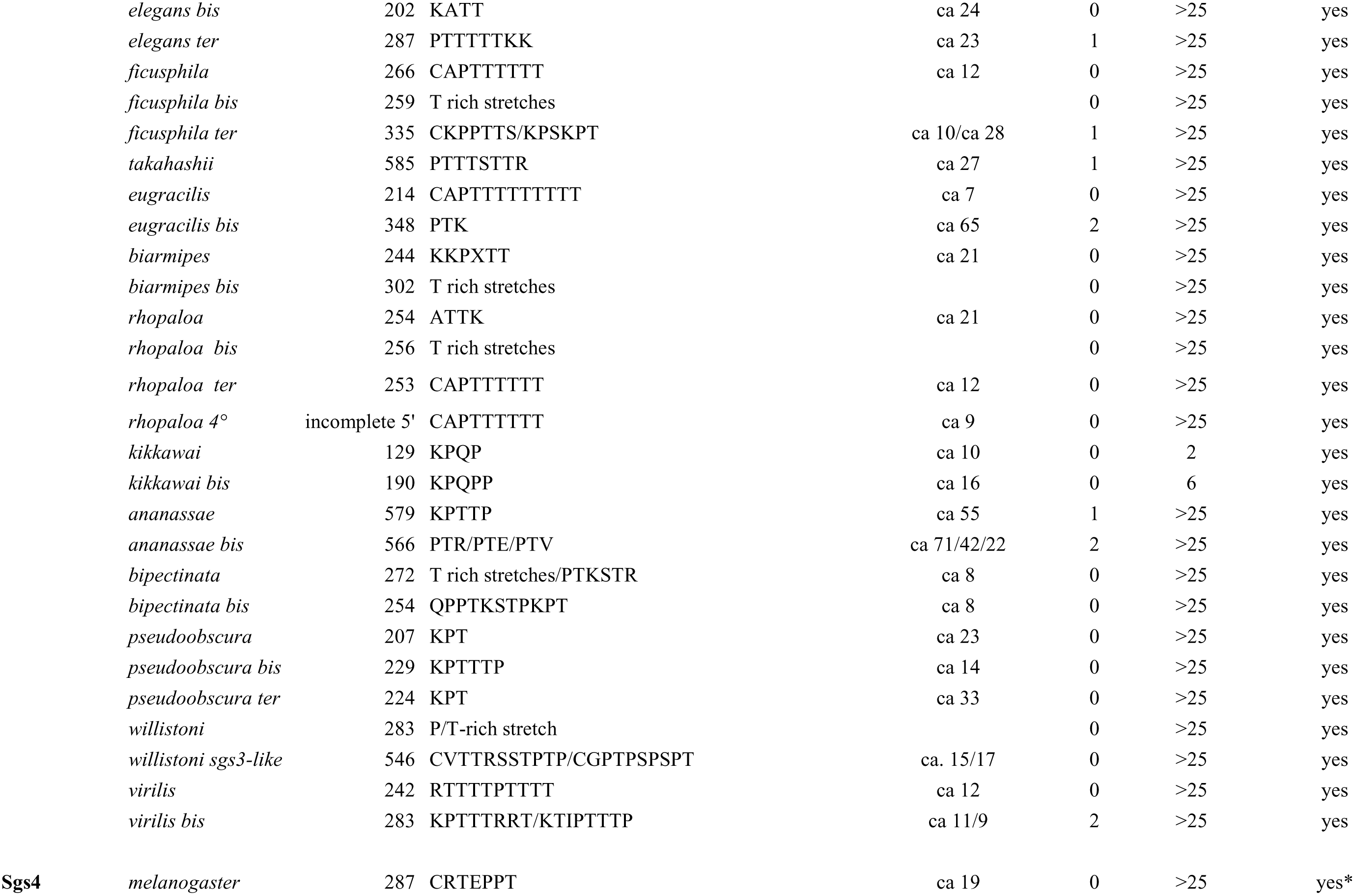
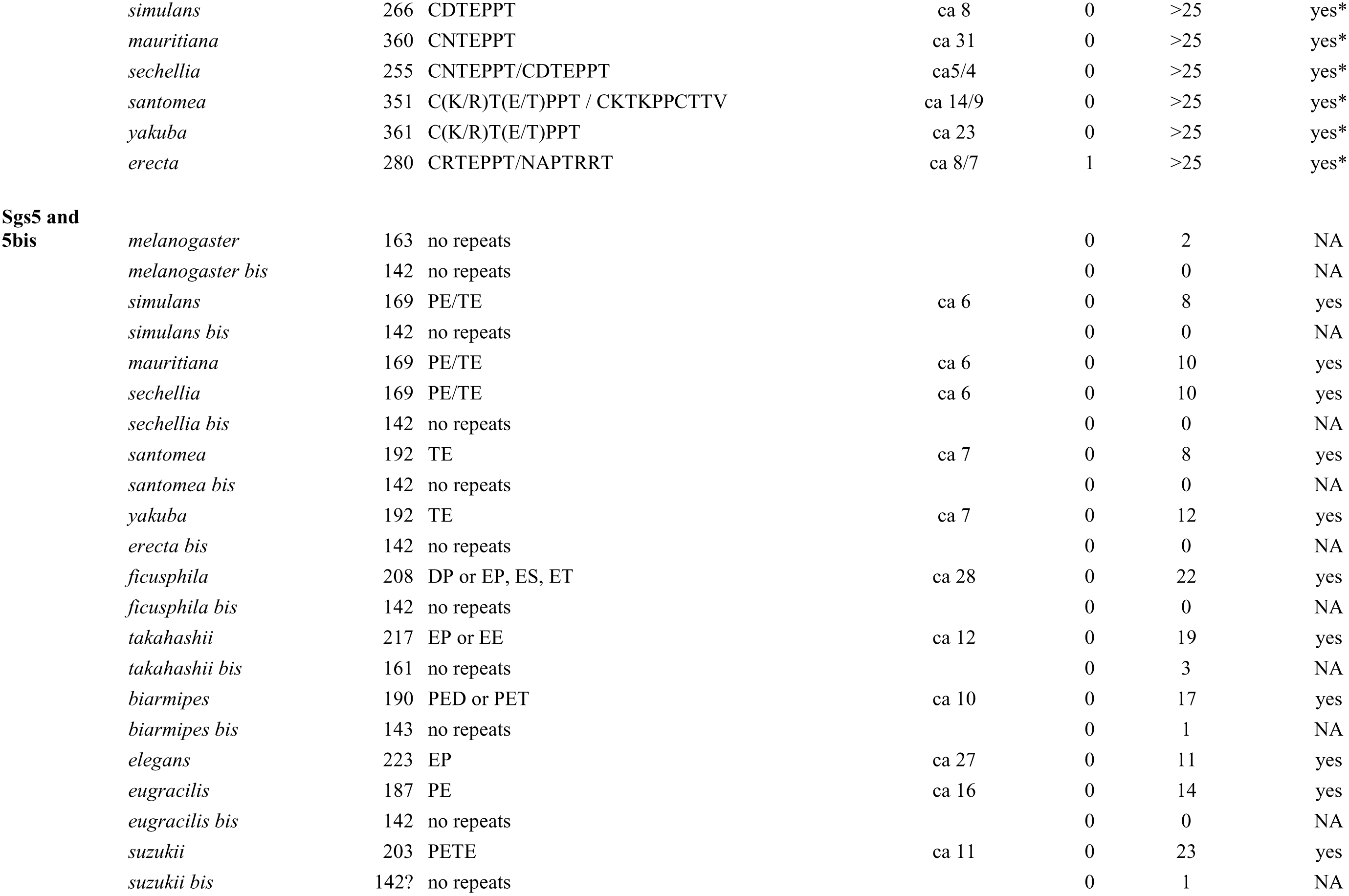
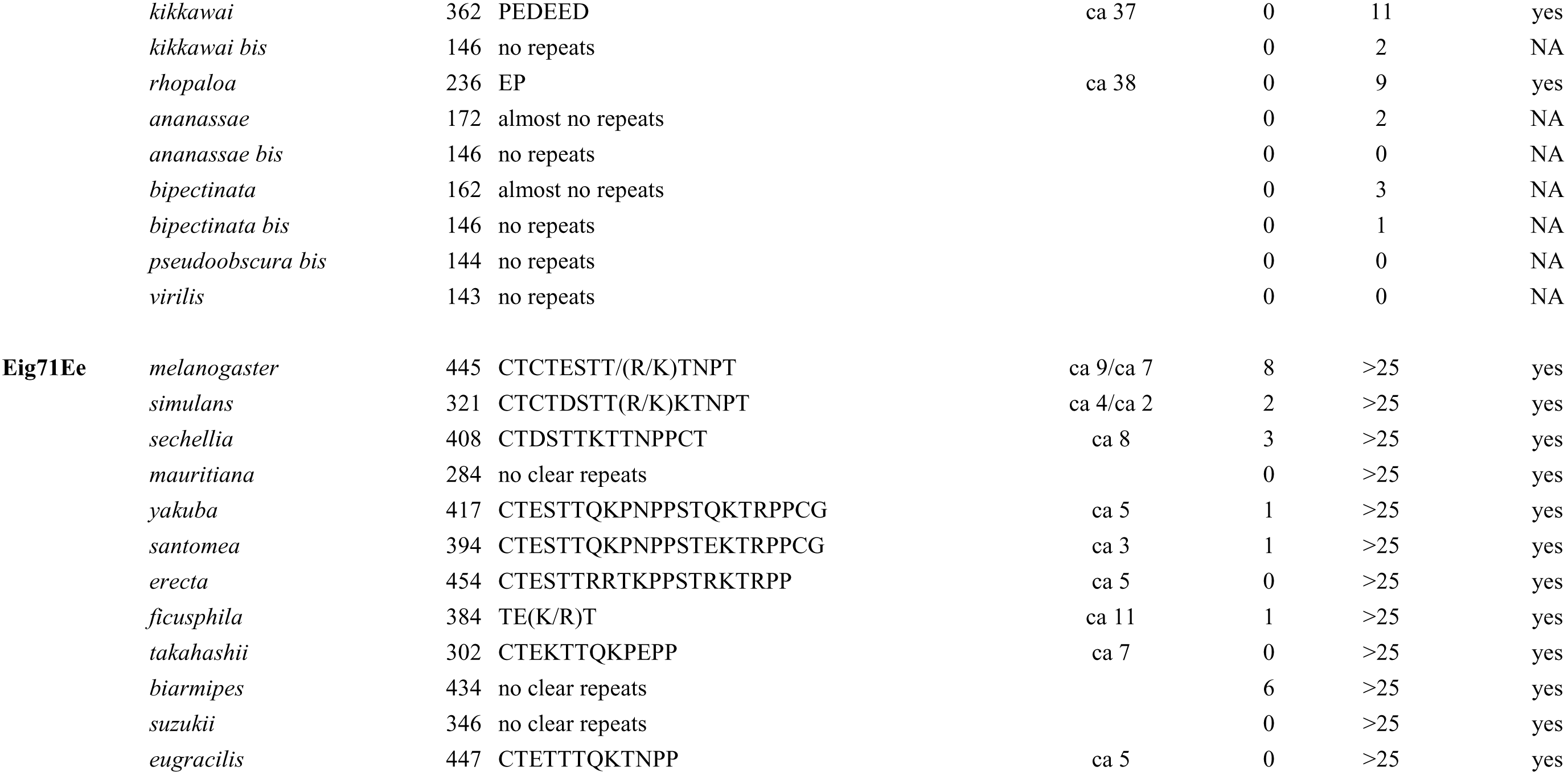
Characteristics of glue proteins in the species studied (except Sgs7 and Sgs8). Glycosylation sites were predicted from http://www.cbs.dtu.dk/services/NetNGlyc and http://www.cbs.dtu.dk/services/NetOGlyc for N glycosylation and O glycosylation, respectively. *: except for IUPred and PrDOS

### Interspecific variation in number and sequence of repeats

Between closely related species the number of repeats varied enormously and the repeated sequence diverged sometimes rapidly (Table 3). *D. simulans*, *D. sechellia,* and *D. mauritiana* form a clade, which split less than 300,000 years ago [40]. Their *Sgs1* genes harbor the same repeated sequence but the number of repeats ranges from 40 in *D. simulans* to 13 and 22 in *D. mauritiana* and *D. sechellia*, respectively. Sgs3 is very similar in the three species, except in the number of repeats. There are no repeats in *D. simulans,* but threonine-rich stretches; in the published sequence of *D. mauritiana*, there are three tandem occurrences of CAPPTRPPCTSP(T)_n_; in *D. sechellia*, several CKP(T)_6_ repeats. Sgs4 shows shared repeats C(D/N)TEPPT among these species, with many more repeats in *D. mauritiana*. In contrast, in the sibling species *D. yakuba* and *D. santomea*, which diverged 0.5 million years ago [41, 42], *Sgs3*, *Sgs4* and *Sgs5* harbor the same repeat sequences and the same number of repeats (Table 3). *Sgs4* genes show 91% identity at the protein level with the same 23 repeats; Sgs5 97% identity. Another pair of species worth of interest is *D. suzukii*/*D. biarmipes*, considered to have diverged ca. 7.3 mya [43]. As mentioned above, only Sgs1 and Sgs5 can be compared because *D. suzukii* has lost *Sgs3*, and *Sgs4* is limited to the *melanogaster* subgroup. Despite a longer divergence time than for the previous comparisons, the Sgs1 29 amino acid repeats are similar in the two species but *D. suzukii* has many more repeat units. In the non repeat parts, identity is 69.3 %; Sgs5 is well conserved even in the repeat region, with an overall identity of 76.4 % in amino acids, and 84.8 % in the non-repeat parts. A last pair of related species (despite their belonging to different subgroups) is *D. elegan*s*/D. rhopalo*a. Their divergent time is unknown. We found that their Sgs proteins are very similar overall, including the repeat parts. This is less striking for the repeats in Sgs3, which exists as four gene copies in *D. rhopalo*a. Their Sgs5 shared a high overall identity (75%), with repeats (Glu-Pro)_n_. In the non-repeat parts, identity rose to 82%. Indeed we often found more divergence among paralogs within a genome than across orthologous proteins.

Structure prediction programs (IUPred [44], PrDOS [45], disEMBL [46], PONDR [47]) indicate that the repeat regions of Sgs1, Sgs3, Sgs4, Sgs5 and Eig71Ee are intrinsically disordered (Fig. 4). Only IUPred and PrDOS indicate Sgs4 repeats to be ordered, in disagreement with the other predictors.

**Figure 4.**
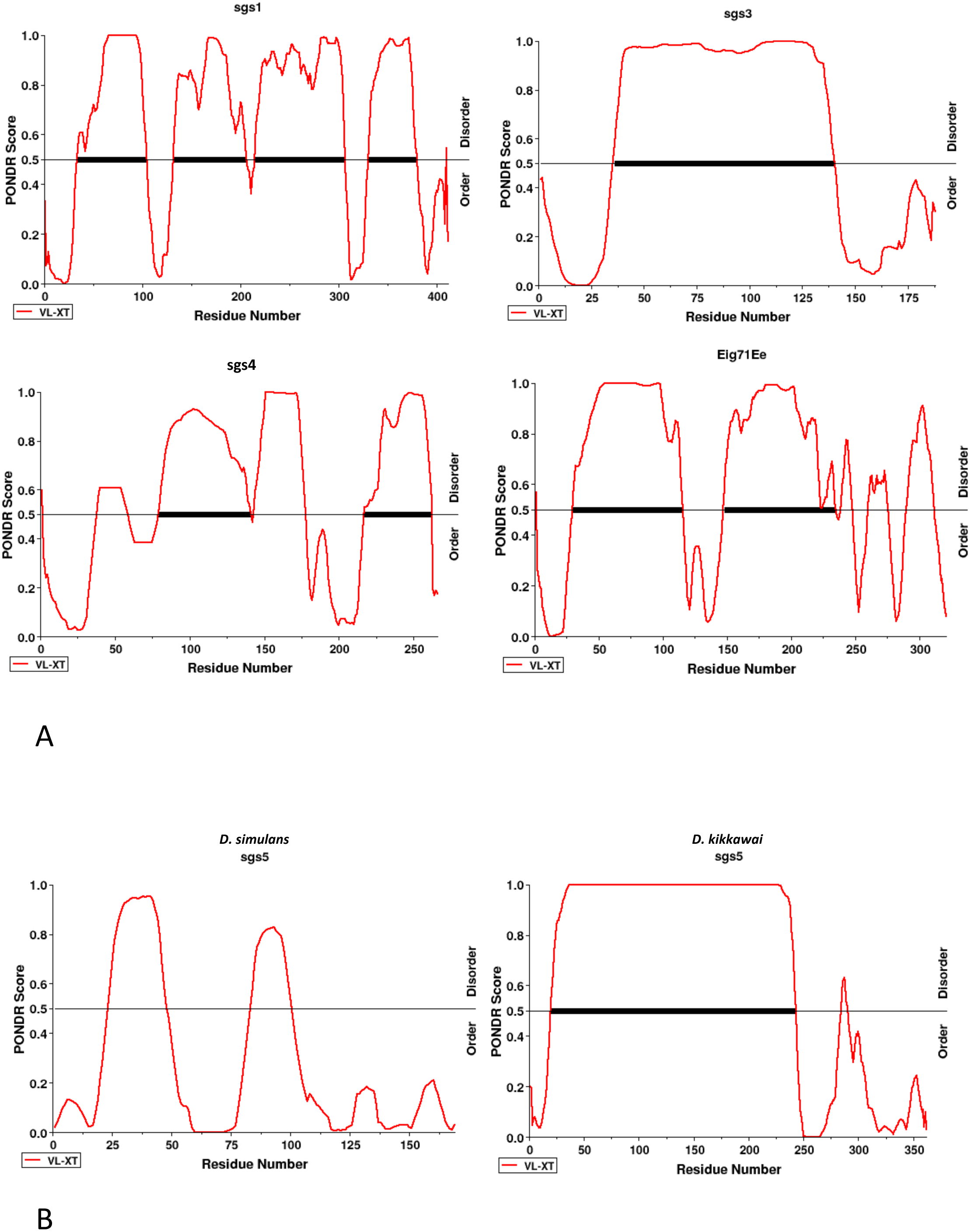
Example of predictions for disordered regions by PONDR. A: The glue proteins with internal repeats of *D. simulans,* except Sgs5, and Eig71Ee; B: example of Sgs5 protein with large internal repeats (*D. kikkawai*) compared to the one of *D. simulans*.

### Intraspecific variation in number of repeats

Owing to the difficulty of short-read sequencing methods to deal with the repeated sequences found in glue genes, we could not get a species-wide insight of repeat number variation (RNV) in *D. melanogaster*. Therefore, we resequenced *Sgs3* and *Sgs4* in strains from various geographic locations using classical Sanger sequencing (Table 4). We found striking inter-and intrapopulation variation in the number of repeats : for *Sgs3* (Fig. S2 and S3, Table 4), there was at least 9 repeat difference between the shortest and the longest allele (22 to 31); for *Sgs4*, 18 to more than 26 repeats (Fig. S4 and S5, Table 4). Regarding the *Sgs4* data from the Drosophila Genome Nexus study (Cairo population), we observed that the repeat region was erroneously reconstituted, often underestimating the repeat number, compared to our Sanger sequencing. We also sequenced the *Sgs3* and *Sgs4* genes in wild-caught *D. mauritiana* individuals. For *Sgs3* we found variation in the number of stretched threonines (10 or 12) and in the number of repeats (Fig. S6A and Table 4). For *Sgs4*, we found that the actual sequences were much longer than the sequence available online, and variable in length, even at the intrapopulation level, ranging from 25 to 35 repeats of the 7 amino acid motif (Fig. S6B and Table 4).

**Table 4:**
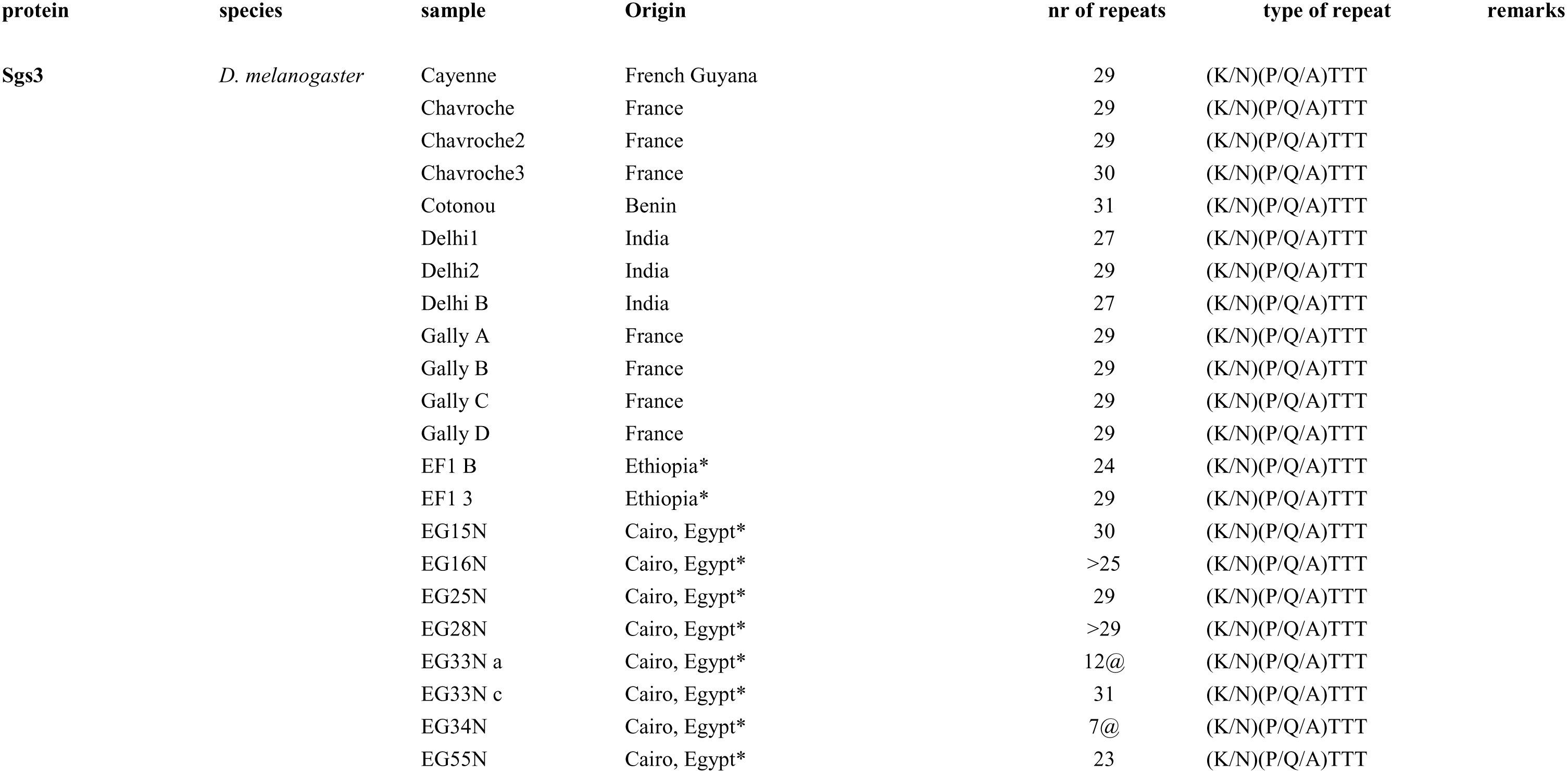
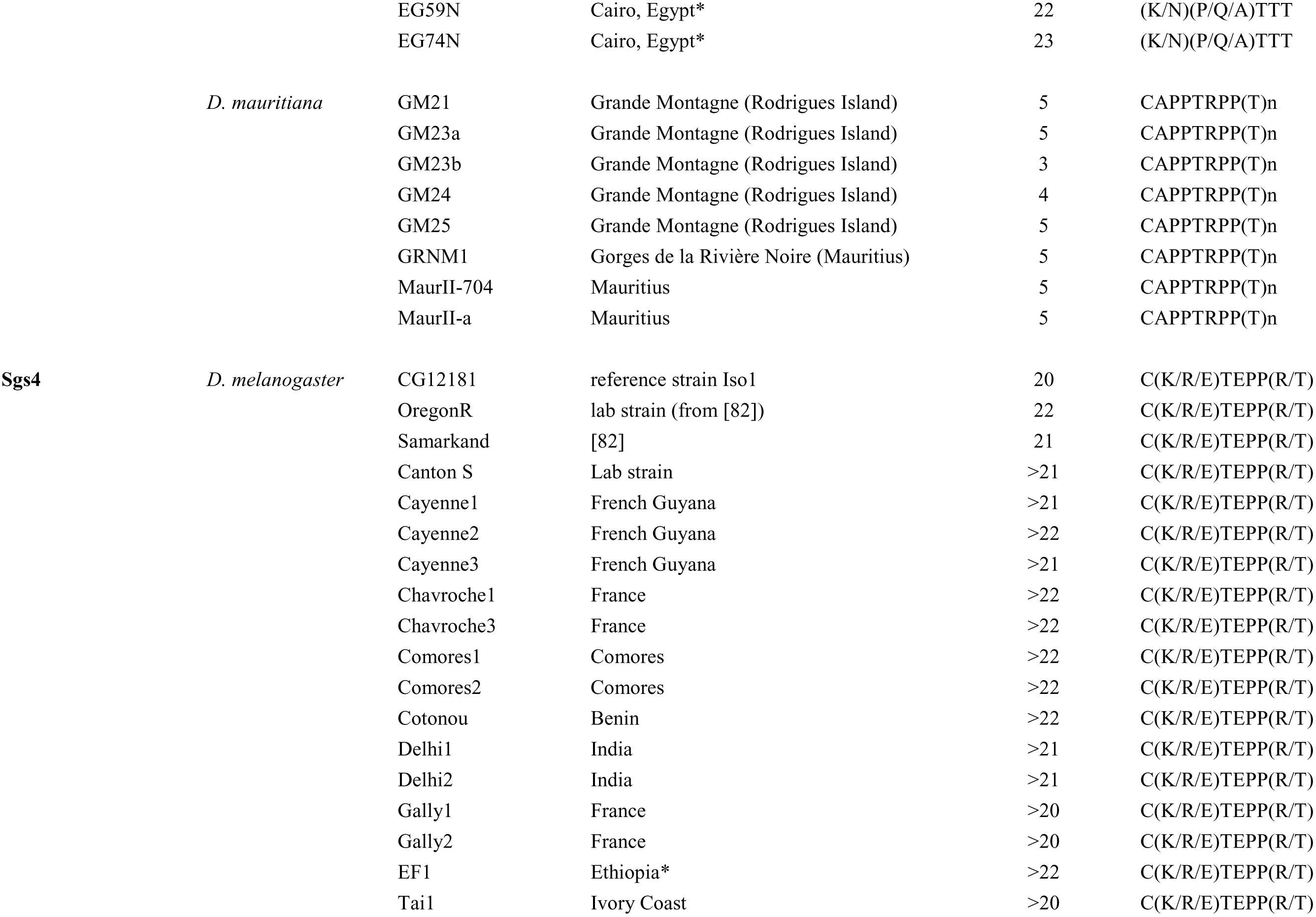
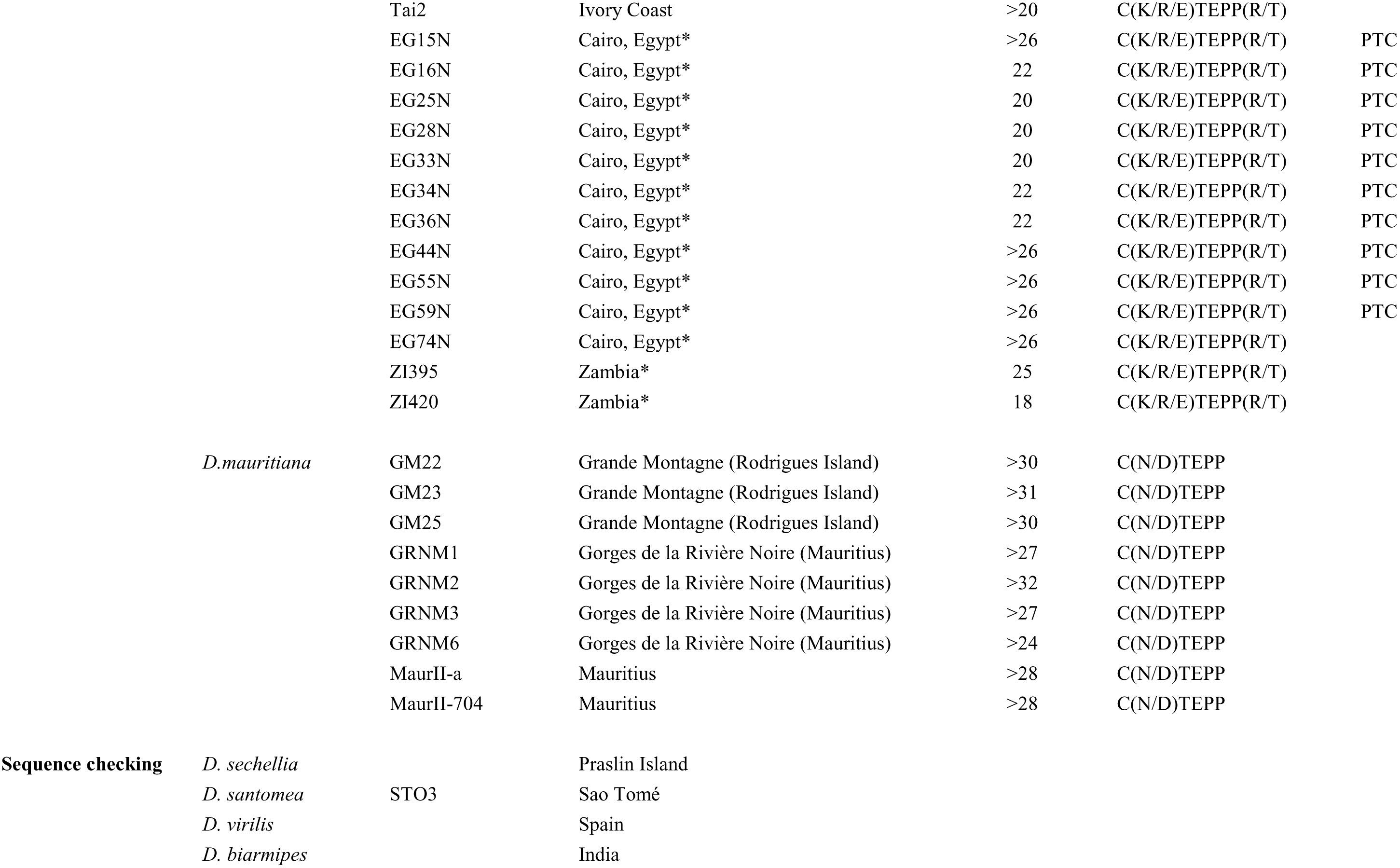
List of strains used for PCR amplification. Number of repeats and repeat motifs in Sgs3 and Sgs4 in populations of *D. melanogaster* and *D. mauritiana*. Sequences of Sgs4 for Oregon R and Samarkand strains are from [82]. * indicate lines also used in the Drosophila Nexus project. @ indicate suspected artifactual repeat losses during cloning. PTC indicates the presence of a premature termination codon.

| protein | species | sample | Origin | nr of repeats | type of repeat | remarks |
| --- | --- | --- | --- | --- | --- | --- |
| Sgs3 | *D. melanogaster* | Cayenne | French Guyana | 29 | (K/N)(P/Q/A)TTT |  |
|  |  | Chavroche | France | 29 | (K/N)(P/Q/A)TTT |  |
|  |  | Chavroche2 | France | 29 | (K/N)(P/Q/A)TTT |  |
|  |  | Chavroche3 | France | 30 | (K/N)(P/Q/A)TTT |  |
|  |  | Cotonou | Benin | 31 | (K/N)(P/Q/A)TTT |  |
|  |  | Delhi1 | India | 27 | (K/N)(P/Q/A)TTT |  |
|  |  | Delhi2 | India | 29 | (K/N)(P/Q/A)TTT |  |
|  |  | Delhi B | India | 27 | (K/N)(P/Q/A)TTT |  |
|  |  | Gally A | France | 29 | (K/N)(P/Q/A)TTT |  |
|  |  | Gally B | France | 29 | (K/N)(P/Q/A)TTT |  |
|  |  | Gally C | France | 29 | (K/N)(P/Q/A)TTT |  |
|  |  | Gally D | France | 29 | (K/N)(P/Q/A)TTT |  |
|  |  | EF1 B | Ethiopia* | 24 | (K/N)(P/Q/A)TTT |  |
|  |  | EF1 3 | Ethiopia* | 29 | (K/N)(P/Q/A)TTT |  |
|  |  | EG15N | Cairo, Egypt* | 30 | (K/N)(P/Q/A)TTT |  |
|  |  | EG16N | Cairo, Egypt* | >25 | (K/N)(P/Q/A)TTT |  |
|  |  | EG25N | Cairo, Egypt* | 29 | (K/N)(P/Q/A)TTT |  |
|  |  | EG28N | Cairo, Egypt* | >29 | (K/N)(P/Q/A)TTT |  |
|  |  | EG33N a | Cairo, Egypt* | 12@ | (K/N)(P/Q/A)TTT |  |
|  |  | EG33N c | Cairo, Egypt* | 31 | (K/N)(P/Q/A)TTT |  |
|  |  | EG34N | Cairo, Egypt* | 7@ | (K/N)(P/Q/A)TTT |  |
|  |  | EG55N | Cairo, Egypt* | 23 | (K/N)(P/Q/A)TTT |  |

|  |  |  |  |  |  |
| --- | --- | --- | --- | --- | --- |
|  |  | EG59N | Cairo, Egypt* | 22 | (K/N)(P/Q/A)TTT |
|  |  | EG74N | Cairo, Egypt* | 23 | (K/N)(P/Q/A)TTT |
|  | *D. mauritiana* | GM21 | Grande Montagne (Rodrigues Island) | 5 | CAPPTRPP(T)n |
|  |  | GM23a | Grande Montagne (Rodrigues Island) | 5 | CAPPTRPP(T)n |
|  |  | GM23b | Grande Montagne (Rodrigues Island) | 3 | CAPPTRPP(T)n |
|  |  | GM24 | Grande Montagne (Rodrigues Island) | 4 | CAPPTRPP(T)n |
|  |  | GM25 | Grande Montagne (Rodrigues Island) | 5 | CAPPTRPP(T)n |
|  |  | GRNM1 | Gorges de la Rivière Noire (Mauritius) | 5 | CAPPTRPP(T)n |
|  |  | MaurII-704 | Mauritius | 5 | CAPPTRPP(T)n |
|  |  | MaurII-a | Mauritius | 5 | CAPPTRPP(T)n |
| **Sgs4** | *D. melanogaster* | CG12181 | reference strain Iso1 | 20 | C(K/R/E)TEPP(R/T) |
|  |  | OregonR | lab strain (from [82]) | 22 | C(K/R/E)TEPP(R/T) |
|  |  | Samarkand | [82] | 21 | C(K/R/E)TEPP(R/T) |
|  |  | Canton S | Lab strain | >21 | C(K/R/E)TEPP(R/T) |
|  |  | Cayenne1 | French Guyana | >21 | C(K/R/E)TEPP(R/T) |
|  |  | Cayenne2 | French Guyana | >22 | C(K/R/E)TEPP(R/T) |
|  |  | Cayenne3 | French Guyana | >21 | C(K/R/E)TEPP(R/T) |
|  |  | Chavroche1 | France | >22 | C(K/R/E)TEPP(R/T) |
|  |  | Chavroche3 | France | >22 | C(K/R/E)TEPP(R/T) |
|  |  | Comores1 | Comores | >22 | C(K/R/E)TEPP(R/T) |
|  |  | Comores2 | Comores | >22 | C(K/R/E)TEPP(R/T) |
|  |  | Cotonou | Benin | >22 | C(K/R/E)TEPP(R/T) |
|  |  | Delhi1 | India | >21 | C(K/R/E)TEPP(R/T) |
|  |  | Delhi2 | India | >21 | C(K/R/E)TEPP(R/T) |
|  |  | Gally1 | France | >20 | C(K/R/E)TEPP(R/T) |
|  |  | Gally2 | France | >20 | C(K/R/E)TEPP(R/T) |
|  |  | EF1 | Ethiopia* | >22 | C(K/R/E)TEPP(R/T) |
|  |  | Tai1 | Ivory Coast | >20 | C(K/R/E)TEPP(R/T) |
|  |  | Tai2 | Ivory Coast | >20 | C(K/R/E)TEPP(R/T) |  |
|  |  | EG15N | Cairo, Egypt* | >26 | C(K/R/E)TEPP(R/T) | PTC |
|  |  | EG16N | Cairo, Egypt* | 22 | C(K/R/E)TEPP(R/T) | PTC |
|  |  | EG25N | Cairo, Egypt* | 20 | C(K/R/E)TEPP(R/T) | PTC |
|  |  | EG28N | Cairo, Egypt* | 20 | C(K/R/E)TEPP(R/T) | PTC |
|  |  | EG33N | Cairo, Egypt* | 20 | C(K/R/E)TEPP(R/T) | PTC |
|  |  | EG34N | Cairo, Egypt* | 22 | C(K/R/E)TEPP(R/T) | PTC |
|  |  | EG36N | Cairo, Egypt* | 22 | C(K/R/E)TEPP(R/T) | PTC |
|  |  | EG44N | Cairo, Egypt* | >26 | C(K/R/E)TEPP(R/T) | PTC |
|  |  | EG55N | Cairo, Egypt* | >26 | C(K/R/E)TEPP(R/T) | PTC |
|  |  | EG59N | Cairo, Egypt* | >26 | C(K/R/E)TEPP(R/T) | PTC |
|  |  | EG74N | Cairo, Egypt* | >26 | C(K/R/E)TEPP(R/T) |  |
|  |  | ZI395 | Zambia* | 25 | C(K/R/E)TEPP(R/T) |  |
|  |  | ZI420 | Zambia* | 18 | C(K/R/E)TEPP(R/T) |  |
|  | *D.mauritiana* | GM22 | Grande Montagne (Rodrigues Island) | >30 | C(N/D)TEPP |  |
|  |  | GM23 | Grande Montagne (Rodrigues Island) | >31 | C(N/D)TEPP |  |
|  |  | GM25 | Grande Montagne (Rodrigues Island) | >30 | C(N/D)TEPP |  |
|  |  | GRNM1 | Gorges de la Rivière Noire (Mauritius) | >27 | C(N/D)TEPP |  |
|  |  | GRNM2 | Gorges de la Rivière Noire (Mauritius) | >32 | C(N/D)TEPP |  |
|  |  | GRNM3 | Gorges de la Rivière Noire (Mauritius) | >27 | C(N/D)TEPP |  |
|  |  | GRNM6 | Gorges de la Rivière Noire (Mauritius) | >24 | C(N/D)TEPP |  |
|  |  | MaurII-a | Mauritius | >28 | C(N/D)TEPP |  |
|  |  | MaurII-704 | Mauritius | >28 | C(N/D)TEPP |  |
| **Sequence checking** | *D. sechellia* |  | Praslin Island |  |  |  |
|  | *D. santomea* | STO3 | Sao Tomé |  |  |  |
|  | *D. virilis* |  | Spain |  |  |  |
|  | *D. biarmipes* |  | India |  |  |  |

### Nonsense mutations in the *Sgs* genes

Despite the rather low quality of sequences in the Drosophila Genome Nexus data set, we searched for putative premature termination codons (PTC) in *Sgs* genes of *D. melanogaster*, which could lead to non-functional proteins. The search was limited to non-repeat regions. We found PTC in *Sgs4* of several lines, that truncated the protein at the beginning of its conserved C-terminal part. We confirmed experimentally the presence of this PTC in 10 lines of the Cairo population EG (K165stop) (Fig. S5 and Table 4). We also found putative PTC for *Sgs5* in a few lines (W161stop, that is sub-terminal, and maybe not detrimental), and experimental verification confirmed it in one Ethiopian line (EF66N); in *Sgs5bis*, we found a putative PTC (C33stop) in six African lines from Rwanda (RG population) and Uganda (UG population). We also found a putative PTC for *Sgs1* in a few lines from USA and Cairo (P49stop), which was confirmed by resequencing the Egyptian line EG36N. This nonsense mutation required two substitutions from CCA to TAA in all cases. Interestingly, EG36N has also a truncated Sgs4. Therefore its glue should be investigated more carefully. In *Sgs3*, no PTC was found. Putative PTC were found for Eig71Ee in two lines, EA90N (S345stop) and RAL894 (W380stop), both in the C-terminal region. One putative PTC was found in *Sgs7* (Q47stop, line USI33), but was not checked experimentally. No PTC was found in *Sgs8* sequences. Stretches of Ns found in non-repeat regions could possibly, at least in some cases, turn out to be true deletions, which deserves further investigation. There is a possibility that some PTCs could experience stop codon readthrough [48] leading to a correct protein, for instance in Sgs4, because the nonsense mutation was not accompanied by other mutations, which would be expected in case of relaxed selection unless the nonsense mutation is very recent. Further studies of the protein content of the salivary glands in those strains will be needed to check whether Sgs4 is produced and if it is full-size.

### Evolutionary rate of *Sgs* protein sequences

Given than glue proteins harbor RNV and given our hypothesis that they could be putative targets for fast selection, we wanted to test whether glue gene coding sequences evolve quickly. To this end, we computed substitution rates of *Sgs* genes between *D. melanogaster* and *D. simulans*, the genomes of which are well annotated. We did not include *Sgs3*, because the internal repeats were very different and not alignable between the two species. This, at any rate, shows that this particular gene evolved rapidly. However, we were able to make an estimate for *Sgs1*, although it had the biggest size and the highest number of repeats, because the repeats were rather similar in *D. melanogaster* and *D. simulans.* We removed the unalignable parts before computation, therefore underestimating the real evolutionary rate. We performed similarly for *Eig71Ee*, *Sgs4* and *Sgs5*. The results are shown on Table 5. We plotted dN and dN/dS for *Sgs* genes on the genome-wide distribution of dN and dN/dS between these species (Fig. 5) using the data of the flyDIVaS database [49]. All dN values were within the highest quartile, and *Sgs1*, *Sgs4* and *Sgs8* were within the highest three centiles. Furthermore, high dN/dS values were found for *Sgs1* (dN/dS=1.393) and *Sgs8* (dN/dS=1.259), indicating accelerated protein evolution. The dN value of *Sgs8* (0.1789) contrasts with the one of its close relative *Sgs7* (0.0475). We wondered if *Sgs8* had also evolved faster than *Sgs7* in other pairs of related species. Table 6 shows the results for other species pairs known to be close relatives : *D. melanogaster/D. sechellia*, *D. simulans/D. sechellia*; *D. yakuba/D. erecta*; *D. biarmipes/D. suzukii*. Whereas the latter two pairs showed no evolutionary rate difference between *Sgs7* and *Sgs8*, comparing *D. simulans* and *D. sechellia* showed a ten times higher dN for *Sgs7* relative to *Sgs8*, a situation opposite to *D. simulans* vs. *D. melanogaster*. In fact, *D. sechellia Sgs7* is more divergent than *D. simulans* from *D. melanogaster Sgs7*, whereas *Sgs8* did not diverged further. Obviously, the small number of substitutions points to a high variance, and the difference may be not significant.

**Figure 5.**
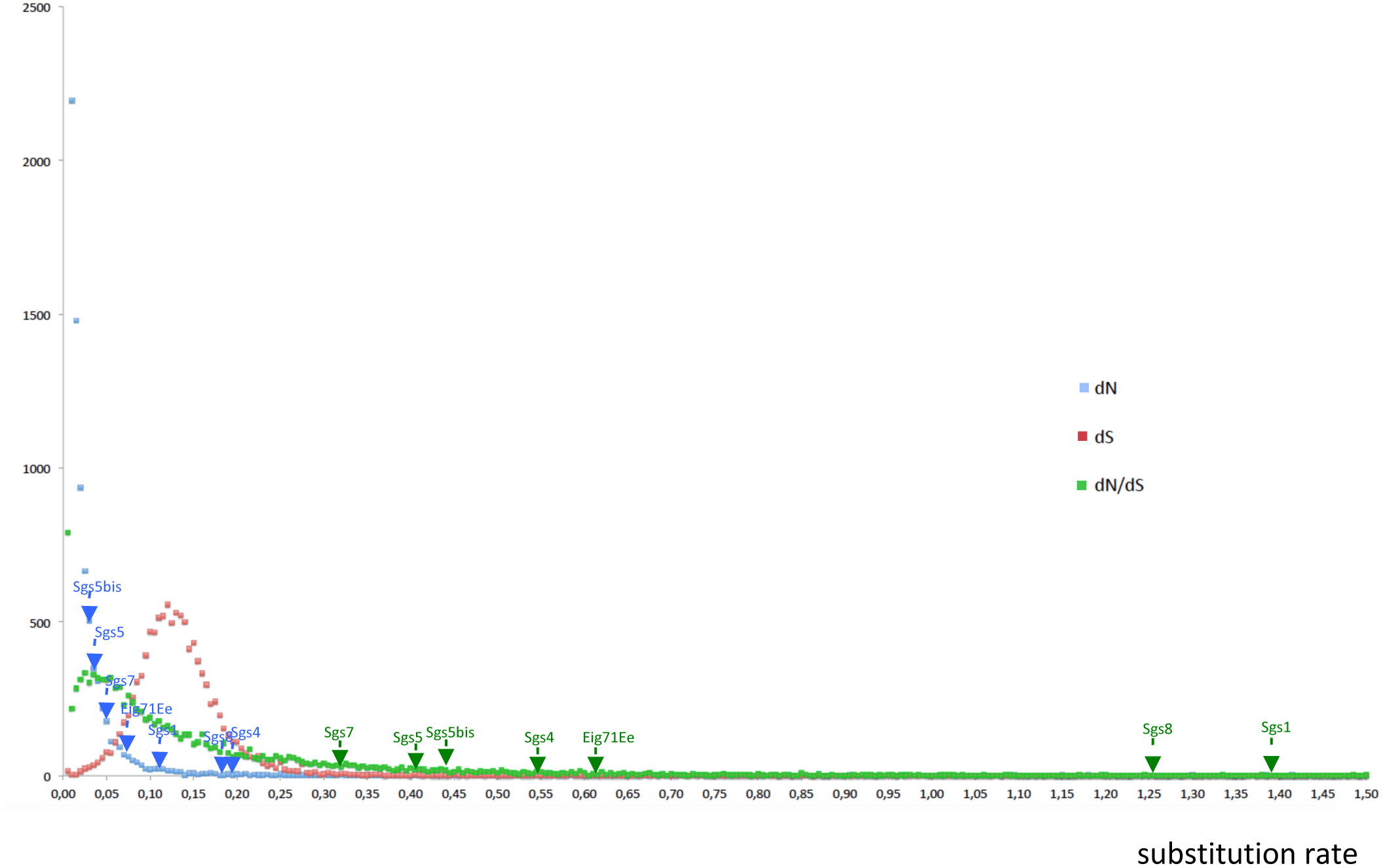
Distribution of dN, dS and dN/dS for the pair *D. melanogaster*/*D. simulans* from the flyDIVaS database with the position of glue genes. Blue dots and arrows : dN; red dots : dS; green dots and arrows: dN/dS. Vertical axis : number of genes. Genes are binned into rate value categories with increment of 0.005.

**Table 5:** Non-synonymous (dN) and synonymous (dS) substitution rates, and the dN/dS ratio for glue genes between *D. melanogaster* and *D. simulans* in pairwise alignments. *Sgs3* was not included, and unalignable regions were removed.

|  | *Sgs1* | *Sgs4* | *Sgs5* | *Sgs5bis* | *Sgs7* | *Sgs8* | *Eig71Ee* |
| --- | --- | --- | --- | --- | --- | --- | --- |
| dN | 0.110 | 0.183 | 0.034 | 0.029 | 0.047 | 0.179 | 0.0678 |
| dS | 0.079 | 0.334 | 0.084 | 0.067 | 0.146 | 0.146 | 0.110 |
| dN/dS | 1.393 | 0.547 | 0.405 | 0.430 | 0.323 | 1.259 | 0.616 |

**Table 6:** Non-synonymous (dN) and synonymous (dS) substitution rates and the ratio dN/dS for *Sgs7* and *Sgs8* between related species pairs in pairwise alignments.

| Species pair | Gene | dN | dS | dN/dS |
| --- | --- | --- | --- | --- |
| *melanogaster/simulans* | *Sgs7* | 0.0475 | 0.1459 | 0.323 |
|  | *Sgs8* | 0.1789 | 0.1420 | 1.259 |
| *melanogaster/sechellia* | *Sgs7* | 0.0990 | 0.1339 | 0.739 |
|  | *Sgs8* | 0.1866 | 0.1216 | 1.534 |
| *simulans/sechellia* | *Sgs7* | 0.0696 | 0.0559 | 1.245 |
|  | *Sgs8* | 0.0060 | 0.0564 | 0.106 |
| *yakuba/erecta* | *Sgs7* | 0.1780 | 0.2235 | 0.796 |
|  | *Sgs8* | 0.1623 | 0.2164 | 0.750 |
| *biarmipes/suzukii* | *Sgs7* | 0.0592 | 0.4329 | 0.137 |
|  | *Sgs8* | 0.0565 | 0.4533 | 0.125 |

To test for adaptive evolution after the “out of Africa” event of *D. melanogaster* [50], we measured the nucleotide diversity π and divergence D_xy_ between one population from Zambia, (ZI) thought to be within the original geographical area of *D. melanogaster*, another African population (EF, Ethiopia) and two derived populations, from France (FR) and USA (Raleigh, RAL). This study was limited to the coding sequences of *Sgs5* and *Sgs5bis*, due to the absence of internal repeats and to the gene size, not too short, (*Sgs7* and *Sgs8* were too short). Due to the numerous residual unidentified nucleotides in the Drosophila Genome Nexus data, the number of sites taken into account could be much smaller than the sequence size, e.g. for *Sgs5bis*, 278 sites left over 489 in RAL. We compared the overall π and D_xy_ between these populations [51]. The results are shown on Table 7. Roughly, for both genes π is higher in ZI than in EF, FR and RAL, as for the whole genome and as expected for the region of origin of this species, but divergences D_xy_ are less than expected from the whole genome, except for the ZI/EF comparison of *Sgs5*. Both genes gave similar results. Therefore, the glue genes *Sgs5* and *Sgs5bis* do not show particular divergence across populations, which could have been related to a change in population environment.

**Table 7:** A) Nucleotide diversity  of *Sgs5* and *Sgs5bis* in four populations, computed from Jukes and Cantor [83] using DnaSP. **B)** Nucleotide divergence between populations D_xy_ computed from Jukes and Cantor in DnaSP. EF: Ethiopia, FR : France, ZI : Zambia, RAL : Raleigh. N : number of lines, n : number of sites, S : number of segregating sites, S.D. : standard deviation, _global_ and D _global_ : nucleotide diversity and nucleotide divergence across the genomes, respectively, from [51].

**A)**
| *Sgs5* | N | n | S | $\pi$ (S.D.) | $\pi_{\text{global}}$ |
| --- | --- | --- | --- | --- | --- |
| **EF** | 35 | 467 | 11 | 0.00450<br>(0.00106) | 0.00622 |
| **FR** | 45 | 476 | 5 | 0.00423<br>(0.00023) | 0.00471 |
| **ZI** | 183 | 489 | 38 | 0.00998<br>(0.00030) | 0.00843 |
| **RAL** | 153 | 386 | 8 | 0.00257<br>(0.00015) | 0.00569 |
| *Sgs5bis* | N | n | S | $\pi$ (S.E.) | $\pi_{\text{global}}$ |
| **EF** | 35 | 406 | 3 | 0.00267<br>(0.00024) | 0.00622 |
| **FR** | 45 | 422 | 8 | 0.00460<br>(0.00029) | 0.00471 |
| **ZI** | 201 | 426 | 37 | 0.00614<br>(0.00034) | 0.00843 |
| **RAL** | 172 | 278 | 5 | 0.00322<br>(0.00018) | 0.00569 |

**B)**
| *Sgs5* | N | n | S | $D_{\square\square}$ (S.D.) | $D_{\square\square \text{ global}}$ |
| --- | --- | --- | --- | --- | --- |
| **ZI/EF** | 183/35 | 467 | 37/11 | 0.01197<br>(0.00082) | 0.00855 |
| **ZI/FR** | 183/45 | 476 | 33/5 | 0.00685<br>(0.00046) | 0.00868 |
| **ZI/RAL** | 183/153 | 386 | 25/8 | 0.00488<br>(0.00036) | 0.00864 |
| **EF/FR** | 35/45 | 454 | 8/5 | 0.00810<br>(0.00128) | 0.00795 |
| **EF/RAL** | 35/153 | 373 | 6/8 | 0.00705<br>(0.00093) | 0.00790 |
| **FR/RAL** | 45/153 | 379 | 7/2 | 0.00162<br>(0.00025) | 0.00546 |

| *Sgs5bis* | **N** | **n** | **S** | **D<sub>□□</sub> (S.D.)** | **D<sub>□□</sub> global** |
| --- | --- | --- | --- | --- | --- |
| **ZI/EF** | 201/35 | 406 | 35/3 | 0.00506<br>(0.00055) | 0.00855 |
| **ZI/FR** | 201/45 | 422 | 36/8 | 0.00639<br>(0.00057) | 0.00868 |
| **ZI/RAL** | 201/172 | 278 | 23/5 | 0.00423<br>(0.00033) | 0.00864 |
| **EF/FR** | 35/45 | 402 | 3/6 | 0.00477<br>(0.00091) | 0.00795 |
| **EF/RAL** | 35/172 | 263 | 3/5 | 0.00551<br>(0.00090) | 0.00790 |
| **FR/RAL** | 45/172 | 276 | 6/5 | 0.00289<br>(0.00035) | 0.00546 |

We also searched for episodic diversifying selection (EDS) among species for the three genes entirely devoid of repeats, *Sgs5bis*, *Sgs7* and *Sgs8*. The branch-site REL test of the HyPhy package was used. No accelerated evolution was detected for *Sgs5bis*, whereas one branch (*D. santomea-D. yakuba* clade) underwent EDS for *Sgs7* (corrected p-value 0.012) and one branch (*D. erecta-D. yakuba-D. santomea)* underwent EDS for *Sgs8* (corrected p-value 0.015) (Fig 6). These results must be considered with caution given the small size of the data set, but anyway do not favor a specific selection regime, regarding single nucleotide (or amino acid) polymorphism.

**Figure 6:**
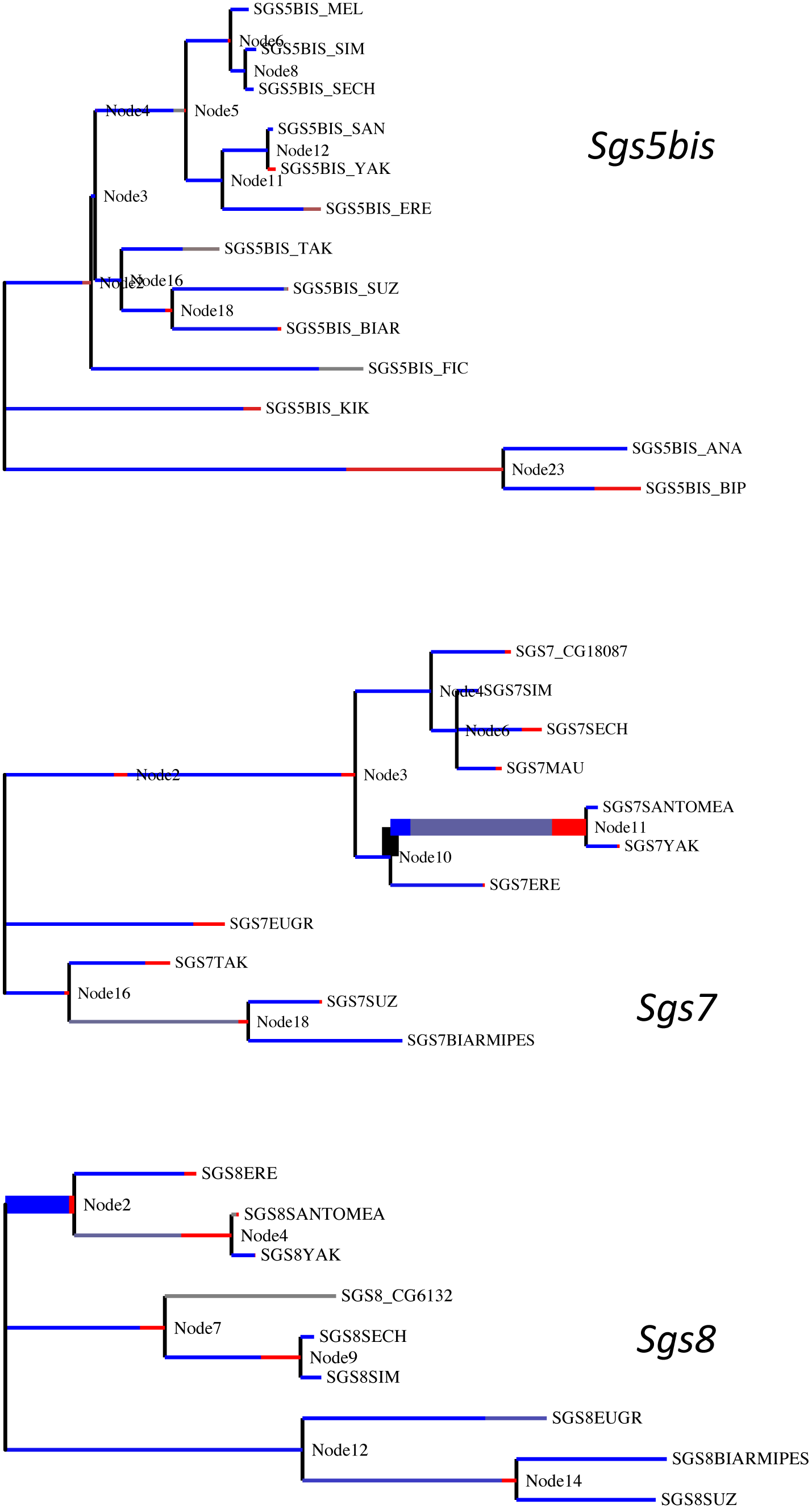
Output trees of BS-REL analyses at Datamonkey website (classic.datamonkey.org). MEL: melanogaster, SIM: simulans, SECH: sechellia, SAN: santomea, YAK: yakuba, ERE: erecta, TAK: takahashii, SUZ: suzukii, BIAR: biarmipes, FIC: ficusphila, KIK: kikkawai, ANA: ananassae, BIP: bipectinata. “The hue of each color indicates strength of selection, with primary red corresponding to    primary blue to w = 0 and grey to w = 1. The width of each color component represent the proportion of sites in the corresponding class. Thicker branches have been classified as undergoing episodic diversifying selection by the sequential likelihood ratio test at corrected p ≤ 0.05”.

## Discussion and conclusion

We have investigated the presence and characteristics of *Sgs* genes and proteins in several *Drosophila* species belonging to the two main subgenera *Sophophora* and *Drosophila*, with particular emphasis on species closer to *D. melanogaster*. We have identified the various *Sgs* genes through sequence similarity with *D. melanogaster*. While this study is extensive, it is of course possible that we may have missed glue genes completely different from the ones of *D. melanogaster*. In order to get the full collection of glue genes we require transcriptional evidence from late larval salivary gland RNA for each species studied. Interestingly, according to our census, the seven genes characterized for years in *D. melanogaster* are far from being always present in the other genomes, although the seven members are generally preserved in the *D. melanogaster* subgroup. Our results are in disagreement with the succinct interspecific study of Farkaš [15]. We also propose here an eighth glue gene, *Sgs5bis.* Based on its close sequence homology and its co-expression with *Sgs5* we propose that these two genes are tandem paralogs. We notice that *Sgs5bis* never contains internal repeats whereas *Sgs5* often harbors more or less developed repeat motifs, although not in *D. melanogaster*. Given our data, and notwithstanding the unbalanced taxonomic sampling which may mislead us, we suggest that the ancestor of the species studied here had only *Sgs3* and *Sgs5bis* (Fig 1). It is likely that *Sgs7*, *Sgs8*, and perhaps also *Sgs1* and *Eig71Ee*, originated from duplications of *Sgs3*. The important differences in repeat motifs between duplicate *Sgs3* (e.g. in *D. eugracilis*) are striking and suggest a high rate of evolution, or independent acquisition of repeats from a repeatless or repeat-poor parental gene. A part of the sequence we named *Sgs3-like* in *D. willistoni* is reported in FlyBase as GK28127, with transcription on the opposite strand, and without a homolog in *D. melanogaster*. Thus, it is possible that some duplicates of *Sgs3* may have been actually recruited for other functions other than glue production. In this respect, it is also possible that Eig71Ee, which has been studied mostly for its immune functions, could be an ancient glue protein, which gained new functions.

The repeat-containing glue proteins are typical of secreted mucins. Mucins are highly glycosylated proteins found in animal mucus and they protect epithelia from physical damage and pathogens [52]. In *D. melanogaster*, more than 30 mucin-like proteins have been identified [53] but the precise function of most of them remain unknown. It would be interesting to compare the glue genes with the other mucin-like genes in terms of protein domains and sequence evolution. In *D. melanogaster*, repeats similar to those of Sgs3 (KPTT) are found in the mucin gene *Muc12Ea*. The high level of glycosylation is thought to favor solubility at high concentration while accumulating in salivary glands ([15]). The richness in cysteines suggests that, upon release in the environment through expectoration, disulfide bridges between glue proteins may be formed by cysteine oxidation by air, making a complex fibrous matrix. Intramolecular disulfide bonds can also be predicted ([15]). Examination of the amino acid composition of the glue proteins suggests that the numerous prolines may induce a zigzag-like shape while serine and threonine, which are very abundant, besides being prone to O-glycosylation, make them very hydrophilic and favor interaction with the solvent and then solubility while preventing folding. The presence of regularly scattered arginines or lysines (or sometimes aspartic and glutamic acids) would add charge repulsion, helping the thread structure to be maintained flat and extended. This is similar to linkers found between mobile domains in some proteins [54]. The shorter Sgs7/Sgs8 would, considering their richness in cysteine, bind the threads together through disulfide bonding.

In the frame of an intrinsically disordered structure (Fig. 4), it is not surprising to observe a high level of repeat number variation (RNV) even at the intra-population level. It has been reported ([55, 56]) that in proteins with internal domain or motif repeats, if these repeats form disordered regions and do not interact with the rest of the protein chain (for a cooperative folding for example), they are more prone to indels which are better tolerated, and favored by the genetic instability of repeated sequences. It is likely that, within a certain repeat number range, variations in repeat numbers might have little effect on the chemical and mechanical properties of the glue. In fact it is likely that the differences in repeat motif sequences rather than the number of repeats would change the mechanical and physical properties of the glue. Accordingly, we measured rather fast rates of evolution, but found no clear indication of positive selection. One reason why the evolution of the repeats is fast (across related species or across paralogs) might be that the constraints to maintain disorder and the thread-like shape are rather loose ([55])

We do not know the respective roles of the different Sgs proteins in the final glue. Farkaš (2016) mentioned that Sgs1 could have chitin-binding properties, which is in line with the function of the glue. He also proposed roles of specific components before expectoration, inside salivary gland granules, related to packaging, solubility… The absence of some glue components may have consequences on its properties and may play a role in adaptation, as suggested by [15]. Gene loss, gene duplication, or repeat sequence change may modify the strength of the glue or its resistance to water or moisture, to acidity (of a fruit) and therefore might be linked to pupariation site preference. *D. suzukii* lacks both Sgs1 and Sgs3, and has duplications of Sgs7. *D. suzukii* pupae are found mostly in the soil just below the surface, and less rarely within ripe and wet fruits such as cherries or raspberries, the pupa half protruding [57, 58]. The extensive loss of *Sgs* genes in *D. suzukii* may be related to its pupariation in soil. Shivanna et al. ([59]) have related pupariation site preference to the quantity of glue and, counter-intuitively, have reported that species that prefer to pupariate on the food medium in the laboratory produce more glue than species that pupariate on the glass walls of the vials. However, the chemical glue content was not investigated. Another study [60] compared pupariation site preferences between the sibling species *D. mauritiana*, *D. sechellia* and *D. simulans*. While *D. simulans* populations from the native region share pupariation preference in fruits with *D. mauritiana* and *D. sechellia*, worldwide populations preferably pupariate off-fruit, i.e. on a drier and harder substrate. Although the QTL associated with pupariation site preference in *D. simulans* and *D. sechellia* do not map to glue genes [60], it would be interesting to see whether, secondarily, significant variations in glue composition or quantity occurred and might be contrasted across *D. simulans* populations. Given its worldwide expansion associated with adaptation to multiple local environments including diverse pupariation sites, *D. melanogaster* is an interesting model to study the intraspecific evolution of *Sgs* genes in relation to adaptation. Interestingly, absence of Sgs4 protein was reported in a few strains from Japan and USA [35], most likely due to deletions or mutations in the promoter region. Our resequencing of a few Nexus lines revealed nonsense mutations within the coding sequence at position 165 in *Sgs4*, deleting the well conserved C-terminal part. The translational consequences for this protein and for final glue properties remain unknown. In addition to such qualitative protein variations, it is possible that the relative proportions of the Sgs proteins in the glue may change in *D. melanogaster* according to the ecological circumstances. In this respect, collecting wandering larvae from various substrates, analyzing their glue composition and designing adhesion assays to compare adhesive properties between various glues will be valuable.

In conclusion, the pupal glue appears as a genetically and phenotypically simple model system for investigating the genetic basis of adaptation. The present work provides a first exploration of the evolution of glue genes across *Drosophila* species and paves the way for future studies on the functional and adaptive consequences of glue composition variation in relation to habitat and geographic and climatic origin.

## Methods

### *Identification of* Sgs *genes in Drosophila species*

The seven annotated glue genes of *D. melanogaster (Sgs1* (CG3047) *; Sgs3* (CG11720) *; Sgs4* (CG12181) *; Sgs5* (CG7596) *; Sgs7* (CG18087) *; Sgs8* (CG6132)) and *Eig71Ee* (CG7604) were used as BLAST queries for retrieving their orthologs in 19 other *Drosophila* species. The genome data used for each species is indicated in Table 1. BLAST searches were performed directly through GenBank, FlyBase [61], the SpottedWingFly base for *D. suzukii* [62] or using local BLAST program (v2.2.25) after downloading the genomes for *D. santomea* [63] and *D. mauritiana* [64]. The BLASTP and TBLASTN programs were used [65], without filtering for low complexity, which otherwise would have missed the repeated regions. Repeats, when present, were often quite different from the repeats present in *D. melanogaster Sgs* sequences. Consequently, BLAST results were often limited to the C-terminal part of the targeted gene, which was the most conserved part of the proteins, and to a lesser extent to the N-terminal end. For each species, a nucleotide sequence containing large regions upstream and downstream of the BLAST hits was downloaded from InsectBase [66] or from species-specific websites when genome data was not present in InsectBase (Table 1). We used Geneious (Biomatters Ltd.) to identify by eye the coding regions, the start of which was identified by the signal peptide sequence. Putative introns were also identified manually, guided by the intron-exon structure of the *D. melanogaster* orthologs. In cases of uncertainties or missing sequence data, we extracted DNA from single flies of the relevant species (Table 4) and the questionable gene regions were amplified with primers chosen in the reliable sequence parts (Table S2), and sequenced by the Sanger method using an ABI 3130 sequencer. For instance, we characterized the exact sequence corresponding to N stretches in the published sequence of *D. mauritiana Sgs4*; we found that the published premature termination codon (PTC) of *D. biarmipes Sgs3* was an error and that three frameshifts found within 50 bp in *D. sechellia Sgs1* were erroneous.

### Evolutionary relationships between genes and estimate of evolutionary rates

Alignments of DNA or protein sequences were done using MUSCLE [67] implemented in Geneious and protein trees were computed using PhyML, as implemented in the online server Phylogeny.fr [68] and drawn using iTOL [69]. The substitution rates dN and dS values for over 10,000 coding sequences computed for *D. melanogaster/D. simulans* comparisons were retrieved from the flyDIVaS database [49] but *Sgs* genes were not included in this dataset. Thus, dN and dS were computed using yn00 in the PAML package ([70]), removing the unalignable parts. We tested for episodic diversifying selection across species using the branch-site random effect likelihood (BS-REL) algorithm implemented in the HyPhy package [71, 72] at the Datamonkey website (classic.datamonkey.org) [73]. We used only genes devoid of repeats to ensure reliable aligments, and we supplied species trees for the analysis.

### Test for accelerated gene turnover

To infer ancestral gene counts in the three newly classified *Sgs* gene families and to determine whether the three newly classified *Sgs* gene families are evolving rapidly we first need to determine the average rate of gene gain and loss (λ) throughout *Drosophila*. Previous studies have estimated λ from 12 *Drosophila* genomes and found rates of 0.0012 gain/losses per million years [4] and 0.006 gains/losses per million years after correcting for assembly and annotation errors [39]. However, since those studies numerous additional *Drosophila* genomes have been published. In order to update the gene gain/loss rate (λ) for this genus, we obtained 25 available *Drosophila* peptide gene annotations from NCBI and FlyBase. The latest versions at the time of study for the genomes of the original 12 sequenced species (*ananassae* v1.05*, erecta* v1.05*, grimshawi* v1.3*, melanogaster* v6.10*, mojavensis* v1.04*, persimilis* v1.3*, pseudoobscura* v3.04*, sechellia* v1.3*, simulans* v2.02*, virilis* v1.06*, willistoni* v1.05*i,* and *yakuba* v1.05) were downloaded from FlyBase [74] and 13 other species (*arizonae, biarmipes, bipectinata, busckii, elegans, eugracilis, ficusphila, kikkawai, miranda, navojoa, rhopaloa, suzukii,* and *takahashii*) were downloaded from NCBI [75].

To ensure that each gene from the 25 *Drosophila* species was counted only once in our gene family analysis, we used only the longest isoform of each protein in each species. We then performed an all-vs-all BLAST search [76] on these filtered sequences. The resulting evalues from the search were used as the main clustering criterion for the MCL (Markov cluster algorithm) program to group peptides into gene families [77].This resulted in 17,330 clusters. We then removed all clusters not present in the *Drosophila* ancestor, resulting in 9,379 gene families. An ultrametric phylogeny with branch lengths in millions of years was inferred using MCL orthogroups in a similar fashion, with the addition of the genome of the house fly, *Musca domestica*, as an outgroup and utilizing single-copy orthogroups between all 26 species [78].

With the gene family data and ultrametric phylogeny as input, we estimated gene gain and loss rates (*λ*) with CAFE v3.0 [4]. This version of CAFE is able to estimate the amount of assembly and annotation error (*ε*) present in the input data using a distribution across the observed gene family counts and a pseudo-likelihood search. CAFE is then able to correct for this error and obtain a more accurate estimate of λ. We find an *ε* of about 0.04, which implies that 4% of gene families have observed counts that are not equal to their true counts. After correcting for this error rate, we find λ = 0.0034. This value for *ε* is on par with those previously reported for *Drosophila* (Table S3; [39]). However, this λ estimate is much higher than the previous reported from 12 *Drosophila* species (Table S3; [4, 39]), indicating a much higher rate of error distributed in such a way that CAFE was unable to correct for it, or a much higher rate of gene family evolution across Drosophila than previously estimated. The 25 species *Drosophila* phylogeny was then manually pruned and modified to represent the 20 *Drosophila* species in which *Sgs* gene families have been annotated. Some *Sgs* gene families are not present in the ancestor of all 20 species, so additional pruning was done to the phylogeny for each family as necessary (see Table S1). The phylogeny, *Sgs* gene copy numbers, and the updated rate of gene gain/loss (λ = 0.0034) were then used by CAFE to infer p-values in each lineage of each family (Table S4). Low p-values (< 0.01) may indicate a greater extent of gene family change along a lineage than is expected with the given λ value, and therefore may represent rapid evolution.

*Search for polymorphism and repeat number variation in* D. melanogaster *and* D. mauritiana Polymorphism in *D. melanogaster* was investigated in the coding regions, especially the repeat number variation (RNV). We intended to use the data from the Drosophila Genome Nexus study ([51, 79], available at the Popfly web site [80]) to assess RNV. This database contains resequenced and aligned genomes of hundreds of *D. melanogaster* lines from about 30 populations from all over the world. Those data, like most *D. melanogaster* populations’ and other species’ genomes were obtained using NGS technologies, which yielded short reads. The data were often not accurate in repeat regions, likely because short reads may be not properly assembled when there are numerous short tandem repeats, and thus could not be used for counting RNV. Thus, experimentally, using single-fly DNAs, we amplified and sequenced the repeat-containing *Sgs3* and *Sgs4* from one or a few individual flies from several strains or natural populations available at the laboratory (French Guyana, Ethiopia, France, Benin, Ivory Coast, India, Comores, and the laboratory strain Canton S), and from a number of lines used in the Drosophila Genome Nexus study (Table 4). In addition, we investigated the occurrence of possible premature termination codons in gene alignments from the Drosophila Nexus database [51, 79], available at the Popfly web site [80] and checked the results by PCR in *Sgs4* and *Sgs5* (Table 4). We also used data from the Drosophila Nexus database to study polymorphism and divergence in *Sgs5* and *Sgs5bis*, which are devoid of repeats, and are not too short. Four populations represented by numerous lines were retained for analysis : ZI (Siavonga, Zambia), for the ancestral geographical range, EF (Fiche, Ethiopia), which shows overall rather large differentiation (Fst) with most other populations [51], and FR (France) and RAL (Raleigh, USA) for the worldwide populations. Diversity and divergence indices were computed with DnaSP [81]. Experimental sequences were deposited to GenBank with accessions MH019984-MH020055.

## Declarations

### Ethics approval and consent to participate

Not applicable

### Consent for publication

Not applicable

### Availability of data and material

Available upon request to the authors

### Authors’ contributions

VCO and JLDL designed the study and analyzed data; JLDL and MB performed experimental work; GWCT performed CAFE analysis; JLDL, VCO and GWCT wrote the manuscript. All authors have read and approved the manuscript.

## Acknowledgments

The authors thank Dr Georges Feller for comments on the disordered protein regions, and Dr Amir Yassin for critical reading of the manuscript. The authors are grateful to three anonymous reviewers for their fruitful comments.

---

**Figure S1:** Ancestral states for the *Sgs1-3-7-8* gene family inferred by CAFE. Species tips are labeled with the observed gene count and internal nodes are labeled with inferred gene counts. Orange branches represent gene losses, blue branches represent gene gains, while black branches represent lineages in which no change in gene copy number is observed. Branches marked with asterisks have marginally significant p-values (< 0.05).

**Figure S2 :** Partial alignment of *Sgs3* sequences with translation in *D. melanogaster* individuals. EF : Ethiopia; Chavroche and Gally : France; Cotonou : Benin; Delhi : India; Cayenne : French Guyana.

**Figure S3:** Partial alignment of *Sgs3* sequences with translation in the EG population (Cairo) of *D. melanogaster*.

**Figure S4:** Partial alignment of *Sgs4* sequences with translation in *D. melanogaster* individuals. EF : Ethiopia; Chavroche and Gally : France; Cotonou : Benin; Delhi : India; Cayenne : French Guyana; Tai : Ivory Coast.

**Figure S5:** Partial alignment of Sgs4 protein sequences in the EG population (Cairo) and ZI (Zambia) of *D. melanogaster*. The reference sequence is shown. Asterisks indicate premature stop codons.

**Figure S6 :** Partial alignment of Sgs3 (A) and Sgs4 (B) amino acid sequences in *D. mauritiana* individuals. Sgs3 mau and Sgs4 mau are the sequences from the online genome. Sgs4 mau has been corrected with our resequencing. Xs are undetermined amino acids.

**Table S1:** Number of gene copies for each family, and results of CAFE analysis for the glue gene families.

**Table S2 :** List of primers used for this study. Different combinations were used to amplify glue genes. All primers were chosen outside the repeated regions. *D. sechellia*, *D. santomea D. virilis* and *D. biarmipes* were resequenced because of uncertainties or putative errors in the online sequences. *D. melanogaster* and *D. mauritiana* were resequenced for studying RNV in *Sgs3* and *Sgs4*.

**Table S3:** Assembly/Annotation error estimation and gene gain/loss rates in a single *λ* model in the 25 *Drosophlia* species included in this study compared to previous studies using fewer species.

**Table S4:** Summary of gene gain and loss events inferred after correcting for annotation and assembly error across all 25 *Drosophila* species. The number of rapidly evolving families is shown in parentheses for each type of change.

